# Molecular Mechanisms of Na^+^-driven Bile Acid Transport in Human NTCP

**DOI:** 10.1101/2023.03.29.534701

**Authors:** Xiaoli Lu, Jing Huang

**Author notes:** Corresponding author: Jing Huang.

## Abstract

Human Na^+^ taurocholate co-transporting protein (hNTCP) is a key bile salt transporter to maintain enterohepatic circulation and is responsible for the recognition of hepatitis B and D viruses (HBV/HDV). Despite recent cryo-EM studies revealing open-pore and inward-facing states of NTCP stabilized by antibodies, the transport mechanism remains largely unknown. To address this knowledge gap, we used molecular dynamics (MD) and enhanced sampling Metadynamics simulations to elucidate the intrinsic mechanism of hNTCP-mediated taurocholate acid (TCA) transport driven by Na^+^-binding. We uncovered three TCA binding modes, including one that closely matched the limited cryo-EM density observed in the open-pore hNTCP. We also captured several key hNTCP conformations in the substrate transport cycle, particularly including an outward-facing, substrate-bound state. Furthermore, we provided thermodynamic evidence supporting that changes in the Na^+^-binding state drive the TCA transport by exploiting the amphiphilic nature of the substrate and modulating the protein environment, thereby enabling the TCA molecule to flip through. Understanding these mechanistic details of Na^+^-driven bile acid transport may aid in the development of hNTCP-targeted therapies for liver diseases.

## Introduction

The human solute carrier 10 (SLC10) protein family encompasses key bile salt transporters that maintain the enterohepatic circulation^1–3^. In this family, the human Na^+^ taurocholate co-transporting protein (hNTCP, encoded by the SLC10A1 gene) is predominantly expressed on the sinusoidal membrane of hepatocytes. It is responsible for the hepatic uptake of bile salts, steroid hormones, thyroid hormones, and various bile acid-conjugated drugs^4–8^. Notably, hNTCP is also the functional cell entry receptor for the hepatitis B and the hepatitis D virus (HBV/HDV)^9^. The viruses use the myristoylated N-terminal preS1 domain (myr-preS1) of the large envelope glycoprotein to recognize and bind to hNTCP^10–13^. Accordingly, hNTCP has emerged as an attractive drug target for developing therapeutics against HBV/HDV infections^14^. To facilitate rational drug discovery, one has to understand not only the recognition mechanisms between hNTCP and HBV but also the intrinsic transport mechanisms of hNTCP to alleviate the potential on-target toxicity.

The cryo-EM structures of NTCP were recently resolved in several studies, in which antibodies or nanobodies were used to stabilize the highly dynamic NTCP^15–19^. In the determined structures, the topology of NTCP consists of the N-terminus on the extracellular surface and nine transmembrane α-helices (TM1-9) in the transmembrane domain (TMD) (Fig. 1a). The TMD is divided into the panel domain (TM1, TM5, and TM6) and the core domain (TM2-4 and TM7-9). In particular, Goutam et al. reported both the inward-facing and the open-pore conformational states of hNTCP, where a major structural rearrangement was observed at the interface between the core and the panel domains^15^. Comparing to inward-facing states, two domains separate from each other on both extracellular and cytoplasmic sides in open-pore hNTCP and open a wide pore throughout the transporter. The amphiphilic substrate taurocholate acid (TCA, Fig. 1b) was co-purified with hNTCP in the experiments, but its coordinates cannot be fully resolved, albeit limited electron density was observed around the residues N103, Q264, and S199 in open-pore hNTCP^15^. The position of the density does not match a previously reported TCA binding pose in the cytoplasmic cavity of an inward-facing conformation of the apical sodium-dependent bile acid transporter (ASBT, also known as SLC10A2), which is a prokaryotic homologue of hNTCP^20^. Two conserved Na^+^-binding sites (Na1: composed of side chains of S105, N106, T123, and E257; Na2: of Q68 and Q261) were inferred to be located at an X-shaped motif (X-motif), which is constructed by the middle-unwound regions of TM3 and TM8. The extracellular surface of the core domain dominates the interaction with antibody fragments, and the transport pathway for substrate and Na^+^ appears to be unimpeded. These structural details provide the structural basis for studying the mechanism of substrate transport.

**Fig 1.**
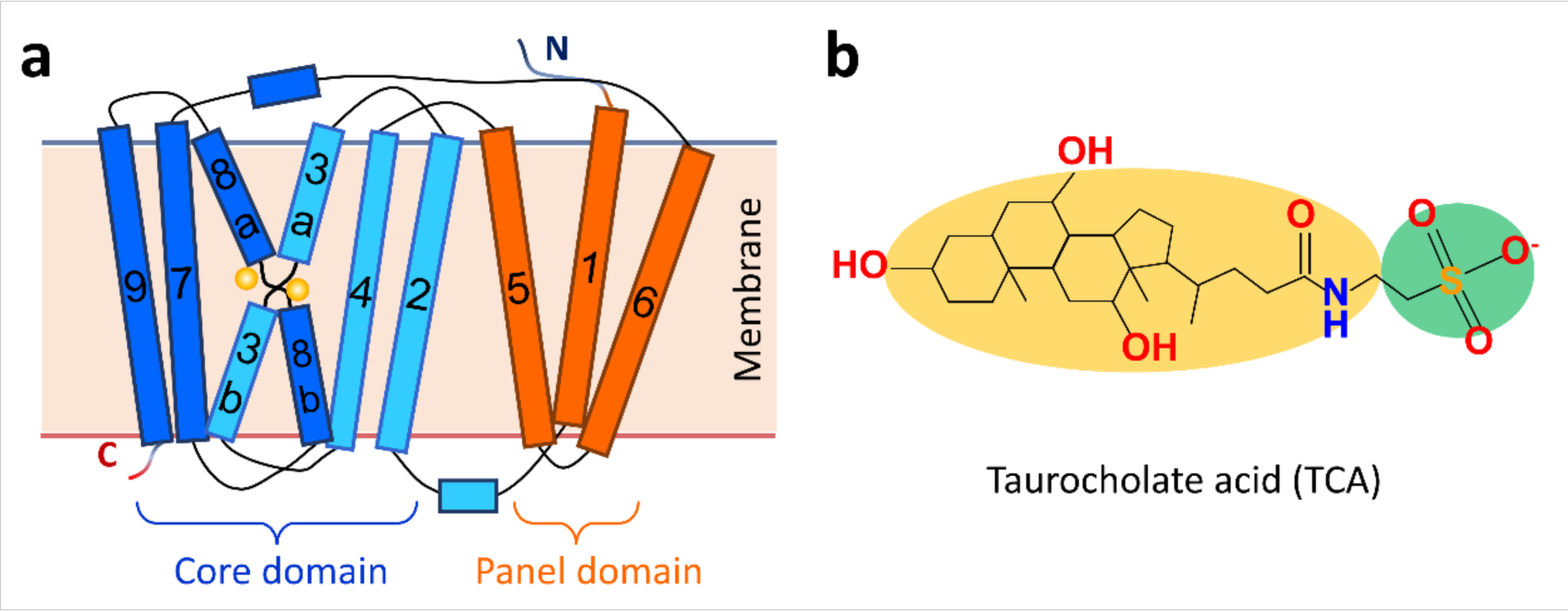
Topology of hNTCP and its substrate TCA. **a** Cartoon representation of NTCP topology. The core domain, panel domain, N-terminus, and C-terminus are labeled. Two Na^+^ are represented as two yellow spheres. **b** Topology of the amphiphilic substrate TCA. The hydrophilic sulfate head group is shaded in green, and the lipophilic sterol tail group is shaded in yellow.

In general, substrate transport by SLC proteins is made possible by large-scale protein conformational changes, especially in the TMD region. A commonly assumed mechanism is the alternating access model^21, 22^, where the transporter alternates between an outward-facing conformation and an inward-facing conformation to allow the substrate to transport. In terms of the details of TMD rearrangement, different kinds of alternating access models have been proposed including rocker-switch, rocking-bundle and elevator-like models^23–26^. The major difference of these transport models lies in the symmetry and rigidity of TM helices when achieving conformational transitions between outward-facing and inward-facing states^27^. However, the experimentally captured open-pore state of hNTCP appears to be inconsistent with the alternating-access transport mechanisms observed in most SLCs, involving additional occluded substrate-bound intermediate of the transport cycle^15^. It’s also long hypothesized that hNTCP exhibits distinctly high dynamics and structural heterogeneity, as hinted in the extreme difficulties in obtaining its static structure^19^.

Furthermore, a mechanistic understanding of the transport cycle of SLCs involves not only the coupling between protein conformational transitions and substrate binding, but also their coupling to the driving force, such as ion concentration gradients^27^. NTCP is a sodium-dependent secondary symporter that employs the Na^+^ gradient across the membrane to drive substrate transport, with a stoichiometry of one substrate to two sodium ions^28, 29^. While it is challenging to establish and maintain a physiological sodium gradient *in vitro*, carefully designed biochemical and biophysical experiments have been carried out to explore the role of Na^+^ in model transporters, such as sodium-coupled amino acid/sugar transporters^30–33^. Many studies have combined detailed structure-function analyses with computer simulations to understand how sodium ions affect substrate binding and release in secondary active Na^+^ transporters, such as Glt_Ph_, LeuT, and SGLT^34–38^. Nevertheless, the specific role of Na^+^ in the transport process of hNTCP remains poorly understood.

In this work, we perform large-scale molecular dynamics (MD) and enhanced sampling metadynamics (MetaD) simulations to reveal the atomistic details of sodium-driven substrate transport in hNTCP. Considering the rate at which hepatocytes uptake bile acid is on the order of minutes^39^, enhanced sampling techniques such as MetaD is necessary to sample the function-relevant conformational ensemble of hNTCP and reconstruct the free-energy surface as a function of selected collective variables (CVs)^40^. By combining extensive long timescale MD simulations and MetaD, we uncover the molecular mechanisms of Na^+^-driven bile acid transport in hNTCP. Several key conformations in the transport cycle are captured, including the outward-bound, inward-bound, and dynamic-apo states. Our simulations also identify three stable TCA-bound conformations and reveal how the binding and transport of TCA are coupled to the Na^+^-binding on the X-shape motif of hNTCP. The corresponding conformational changes are quantified by constructing the free energy profiles for Na^+^-bound and Na^+^-unbound states. Our study provides a thermodynamic explanation on how the ion gradients drive the substrate transport. The atomistic details on the mechanism of Na^+^-gated hNTCP transport substrates will facilitate the development of hNTCP-targeted therapies for liver diseases.

## Results

### Binding modes of substrate TCA in hNTCP

To understand the substrate transport mechanism, we first assess the potential substrate binding modes in the experimentally resolved open-pore and inward-facing conformations of hNTCP (PDB id: 7PQQ and 7PQG). The two conformations differ mainly in the interface between the core and the panel domains. The different TM arrangements lead to a difference of 4 Å, 5 Å, and 6 Å in the size of the extracellular gate, the middle pore, and the intracellular gate, respectively (Fig. 2a). Targeted on these two conformations, the induced-fit docking was performed using pockets defined by all residues within 5 Å of N103, Q264, and S199 in hNTCP.

**Fig 2.**
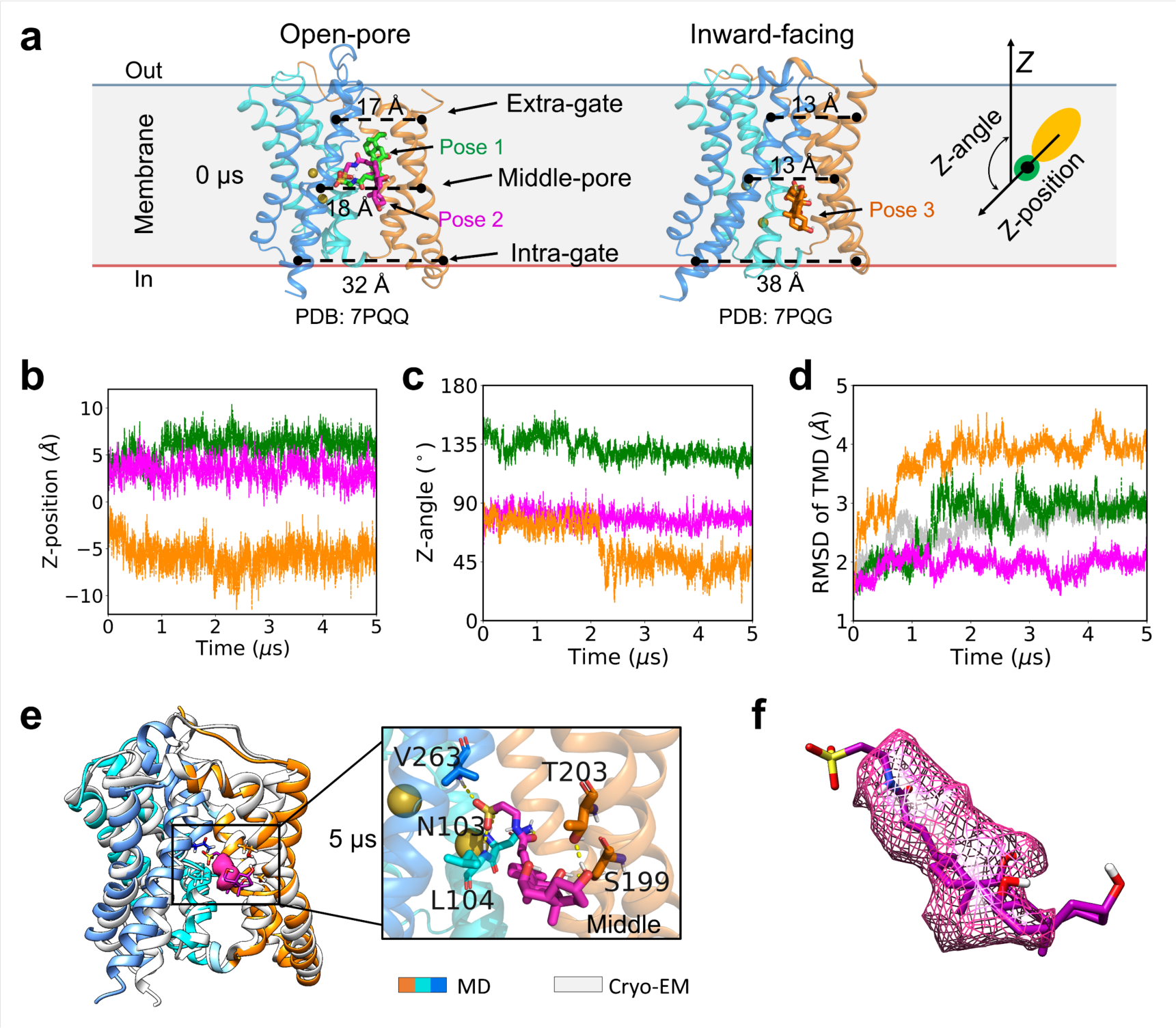
Substrate binding modes of hNTCP. **a** Three initial docking poses in the open-pore and inward-facing conformations of hNTCP, and the cartoon representation of the substrate TCA. **b-d** Time evolution of the Z-Position and Z-angle of TCA, as well as the TMD RMSDs along 5 μs MD simulations of the three TCA-bound hNTCP systems (green: pose 1, magenta: pose 2, orange: pose 3). The TMD RMSD along 5 μs MD simulation of the apo hNTCP system is shown in gray as a reference in panel **d**. **e** Structural alignment of the cryo-EM structure (in grey) and the TCA-bound open-pore hNTCP after 5 μs MD simulation (in colorful). The limited experimental cryo-EM density postulated to be the substrate is marked using magenta mesh. A zoom-in view on the intermolecular interactions between TCA and hNTCP is provided with the residues involved in hydrogen bonding interactions labeled and the two Na^+^ represented as yellow balls. **f** Zoom-in visualization of TCA alignment with the partial cryo-EM density.

For the open-pore hNTCP, molecular docking of the TCA results in two dominant docking poses (poses 1 and 2, green and magenta in Fig. 2a, respectively) with the highest docking scores of −8.00 and −7.99 kcal mol^−1^. The docking poses 1 and 2 share the same configuration of the sulfate head contacting the X-motif (^98^CSPGGNLSN^106^ on TM3 and ^257^ETGCQNVQ^264^ on TM8) while differ in the orientation of the sterol tail. For the inward-facing hNTCP, the molecular docking results in one dominant docking pose with the highest docking scores of −9.04 kcal mol^−1^ (pose 3, orange in Fig. 2a). In the docking pose 3 of inward-facing hNTCP, the sulfate head of TCA is also in contact with the X-motif, while the sterol tail points toward the cytoplasmic side. We note that the docking pose 3 in the intracellular pocket is different from substrate binding pose in the X-ray structure of TCA-bound ASBT (PDB id: 3ZUY and 3ZUX)^20^, in which the sulfate head of TCA is oriented toward the solvent environment and the substrate is closer to the membrane-solvent interface.

We carried out three 5 μs MD simulations, each initialized with one of the three TCA-protein docking structures (Supplementary Table 1, Supplementary Fig. 1). The conformation of TCA in the complex was characterized by the relative position of its sulfate head along the membrane normal (Z-position) with the bilayer midplane at Z = 0 Å, and the angle between its tail-to-head vector and the normal vector of the membrane plane (Z-angle) (Fig. 2a). The time evolution of the Z-position and Z-angle showed that TCA reached to three different equilibrium states based on the initial docking pose (Fig. 2b-c). Starting with the docking pose 1, the Z-position of TCA shifted up to approximately 7 Å and the Z-angle gradually rotates to ∼130° to reach the equilibrium binding pose 1 in the extracellular pocket. Starting with the docking pose 2, the Z-position of TCA remains stable at ∼2 Å and the Z-angle at ∼85° throughout the simulation, maintaining the equilibrium binding pose 2 in the middle pore. Starting with the docking pose 3, the Z-position of TCA shifted down to approximately −5 Å and the Z-angle rotated to ∼45° after 2 μs MD simulation to reach the equilibrium binding pose 3 in the intracellular pocket. Ultimately, three substrate binding sites located respectively in the top extracellular pocket, the middle pore, and the bottom cytoplasmic pocket, were captured with TCA stably bound from 5 μs MD simulations (Supplementary Fig. 2).

Interestingly, the hNTCP protein shows different dynamics in the three TCA-bound states. The initial state with docking pose 1 or 3 and apo state lead the conformational changes larger than ∼3 Å RMSD in the TMD region. By contrast, the TMD exhibits the most significant structural stability when initialized with the docking pose 2, with only less ∼2 Å RMSD change compared to the starting cryo-EM structure (Fig. 2d). Structural alignment confirms the high consistency between the experimentally determined open-pore hNTCP and the TCA-bound hNTCP with binding pose 2 (Fig. 2e). We note that in this state the coordinates of TCA overlap well with the partial cryo-EM density observed at the interface between the core and the panel domain (EMDB id: EMD-13596)^15^. In addition, Liu et al. constructed a complex structure featuring two glyco-chenodeoxycholic acid (GCDC) molecules bound to hNTCP based on two sets of limited cryo-EM density^18^. One of the GCDC molecules occupies the same position as our binding pose 2 (Supplementary Fig. 3a).

While the limited experimental cryo-EM density alone was insufficient to directly model the full substrate-bound complex structure, here a clear binding mode of substrate in the open-pore hNTCP is obtained at the atomic level combining docking and long timescale MD simulations. A close examination on this binding mode shows that residues in the X-motif form multiple hydrogen bonds (H-bonds) with the head group of TCA to maintain its stability. As zooming in Fig. 2e, the backbone -NH groups of N103, L104 and V263 form three H-bonds with the sulfonate group, and the sidechain of N103 forms an additional H-bond with the substrate carbonyl group. Residues on the panel domain (S199 and T203 on TM6) contribute to holding the sterol tail of TCA.

Interestingly, although inserting TCA in the top extracellular pocket or the bottom cytoplasmic pocket of hNTCP induces distinct conformational changes, the complex structures reach equilibrium after approximately 2 μs (green and orange in Fig. 2d). While the TCA head group moves upward (for binding pose 1) or downward (for binding pose 3), it is still anchored by the X-motif through hydrogen bond interactions in either pocket (Supplementary Fig. 4). When TCA binds in the top extracellular pocket, its sterol tail is hold by the residues on the panel domain (S28 on TM1 and N209 on TM6), whereas the tail becomes free when it binds in the bottom cytoplasmic pocket (Supplementary Fig. 4).

Among the multiple binding poses captured, a notable common feature is that the acidic group of the TCA interact directly with the X-motif. This feature can also be identified in the X-ray structures of ASBT bound to citric acid or glycine (PDB id: 4N7W, 6LGV, 6LGY, 6LH1)^41, 42^. In these structures, citric acid interacts directly with the X-motif in the central cavity of ASBT, similar to the sulfate group of TCA in hNTCP (Supplementary Fig. 3b). As for the sterol tail of substrate, the specific binding conformations in the central cavity are seldom reported, potentially due to the instability of sterol binding causing it difficult to be resolve in experimental complex structures. Essentially, secondary active transporters operate as cyclical non-equilibrium systems that couple the spontaneous influx of a driving ion to the flux of the substrate^43^. Multiple binding events can happen along the substrate transport path as these transporters undergo cyclical changes between inward-facing and outward-facing conformations. In hNTCP, the two different electron densities detected by Liu et al. suggest that at least two substrate binding events occurred during the transport process^18^. The X-ray structures of ASBT bound to TCA imply another binding event in the intracellular pocket^20^. Derivation of these intermediate TCA-bound conformational states already reveals the essential process of TCA being transported from the extracellular to the intracellular matrix. Furthermore, the cooperation between substrate binding and synchronized protein conformational changes was captured in microsecond-timescale MD simulations.

### Alternative conformations of hNTCP in TCA-bound and apo states

We further computed the sizes of the extracellular gate, the middle pore, and the intracellular gate of hNTCP along all MD simulations of TCA-bound hNTCP systems. While gauging the extracellular and the intracellular gates is sufficient to distinguish the global conformations for most SLCs such as secondary active major facilitator superfamily (MFS) transporters^44^, for hNTCP additional characterization on the middle pore is found to be indispensable. The size of the middle pore can be used to inspect the main mode of motion in the global conformational changes of hNTCP transport substrate.

The time evolution of the size of extracellular gate, middle pore, and intracellular gate indicates that the global conformation of hNTCP can exhibit the outward-facing, open-pore, and inward-facing states with TCA binding restriction (Fig. 3a-c, Supplementary Fig. 5). When TCA binds to the extracellular pocket, the size of the extracellular gate enlarges to ∼22 Å and the middle pore shrinks to ∼15 Å. Consequently, an additional outward-facing state of hNTCP is sampled with TCA binding to the extracellular pocket. For typical transporters, inward-facing conformations are generally more prevalent, suggesting that outward-facing conformations represent either a higher-energy or a transient state within the transporting cycle, making them more difficult to capture^45^. Interestingly, Park et al. showed that an outward-facing conformation of hNTCP was likely captured under the binding of YN69083 Fab^16^. The RMSDs between this structure (PDB id: 7FCI) and the open-pore and inward-facing structures (7PQQ and 7PQG) are 2.1 Å and 4.3 Å, respectively. The RMSD between the 7FCI structure and our sampled outward-facing structure is 1.4 Å (Supplementary Fig. 6). Examination of the gate sizes revealed that the sampled outward-facing conformation is more open towards the matrix as evidenced by its larger extracellular gate size (∼22 Å compared to ∼17 Å for 7FCI), while they have the similar sizes in the middle pore (∼16 Å and ∼15 Å) and intracellular gate (∼33 Å and ∼32 Å).

**Fig 3.**
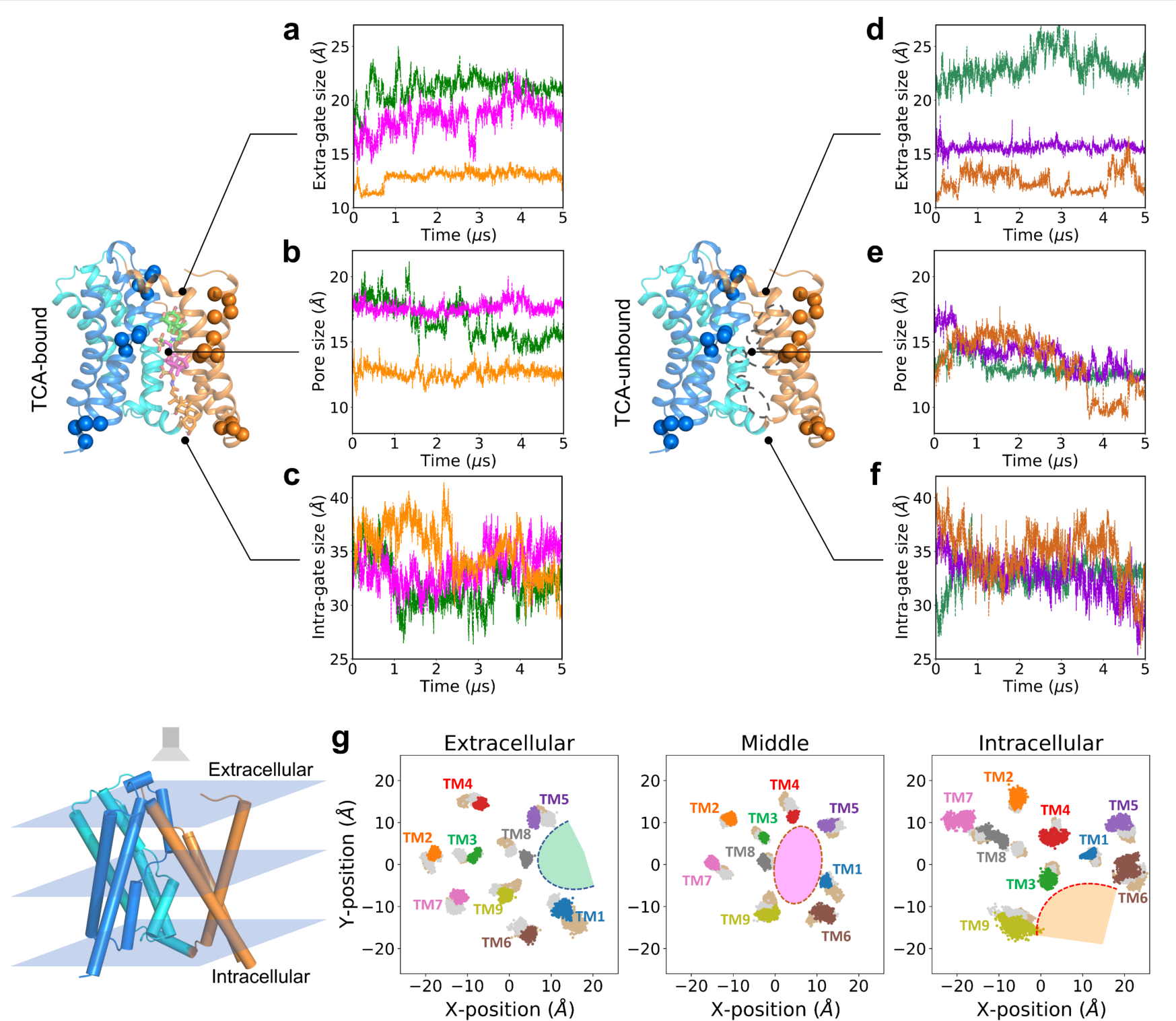
hNTCP exhibits the multiple conformations in the TCA-bound and apo states. **a-f** Time evolution of the size of the extracellular gate, middle pore, and intracellular gate for the three TCA-bound hNTCP (green: outward-facing, magenta: open-pore, orange: inward-facing) and corresponding three apo hNTCP systems after removing TCA from the complexes. **g** The movement of each TM helix in apo hNTCP. The projection of nice TM helices on the membrane-plane depicted from three intersections including the extracellular (left), the middle-membrane (middle), and the intracellular (right) planes. For each helix, the locations sampled starting from the outward-facing state are shown in wheat, the locations sampled starting from the open-pore state are shown in grey, the locations sampled starting from the inward-facing state are shown multicolored.

When TCA binds to the middle pore with binding pose 2, the extracellular gate adopts a smaller size of ∼18 Å and the middle pore remains open for ∼18 Å. Comparing two systems initialized with two different docking poses in open-pore hNTCP, we note that docking pose 1 drives the system out of equilibrium, reaching to an equilibrated outward-facing state. When TCA binds to the cytoplasmic pocket with docking pose 3, the extracellular gate adopts the smallest size of ∼13 Å and the middle pore keeps closed at ∼13 Å. In contrast, the dimension of the intracellular gate shows substantially greater flexibility than the extracellular gate and the middle pore (Fig. 3c). Consequently, no distinct differences emerge among the three TCA-bound states along 5 μs MD simulations.

Three additional 5 μs MD simulations were carried out for apo hNTCP starting from the outward-facing, open-pore, and inward-facing states (Supplementary Fig. 7) after removing both the substrate TCA and two Na^+^ from the complexes. Time evolution of the size of extracellular gate, middle pore, and intracellular gate showed that the extracellular gate was relatively stable, while the middle pore further shrunk to the narrower state, and the intracellular gate remained highly dynamics (Fig. 3d-f). The shrinkage of the middle pore suggests that the instantaneous removal of TCA drives hNTCP out of equilibrium.

To better understand the movement of TMD in apo hNTCP, the projection of nice TM helices on the membrane plane was depicted from three intersections including the extracellular plane, the middle-membrane plane, and the intracellular plane. As shown in Fig. 3g, the projected positions overlapped between the three independent MD simulations of apo hNTCP, suggesting that each TM helix has the flexibility and mobility to access the conformational space sampled by another simulation. Additionally, the extracellular pocket (bounded by TM1, TM5, and TM8) and the intracellular pocket (bounded by TM3, TM6, and TM9) of apo hNTCP are illustrated in Fig. 3g. These two pockets not only have access to the water environment, but also are extensively exposed to the lipid bilayer environment, with phospholipid molecules entering both the extracellular and intracellular pockets, as observed in the MD simulations of apo hNTCP (Supplementary Fig. 8).

The observed multiple conformations of TCA-bound and apo hNTCP provide insights into the substrate transport mechanism of hNTCP. Structural characterization of three TCA-bound hNTCP reveals the high flexibility of the interface between the core and panel domains, which enables the conformational transitions of TMD to cooperate with substrate transport. Two helices, TM6 and TM9, dominantly contribute to the high dynamics of hNTCP through bending and turning respectively, with the fluctuations of ∼15° in angles perpendicular to the membrane (Supplementary Fig. 9). By delineating the solvent-accessible space in both extracellular and intracellular sides, all representative alternative conformations of hNTCP sampled from MD simulations are ordered according to the states of substrate accesses from extracellular side to intracellular side (Fig. 4). First, hNTCP assumes a dynamic-apo state when no substrate access to the extracellular side, and the middle pore is closed. Second, hNTCP adopts an outward-bound state when the substrate enters the extracellular pocket. Next, hNTCP transitions to an open-pore state when the substrate binds to the middle pore. Subsequently, hNTCP manifests an inward-bound state when the substrate binds to the intracellular pocket. Finally, when the substrate completely detaches from the intracellular side, hNTCP reverts to the dynamic-apo state with a closed middle pore.

**Fig. 4.**
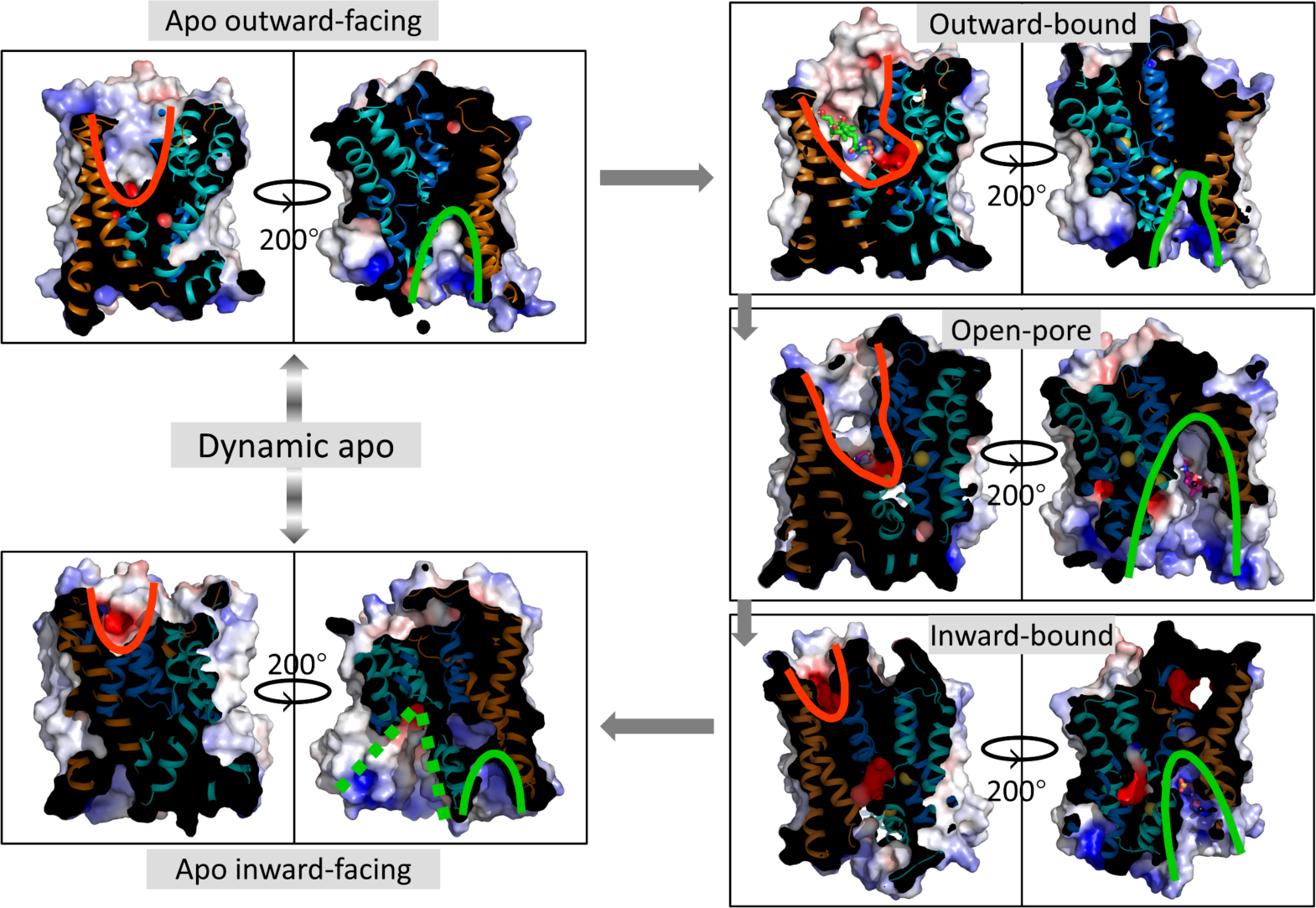
Key conformations of hNTCP sampled from a set of MD simulations for TCA-bound and apo hNTCP. Red and green lines delineate the solvent-accessible space in the both extracellular and intracellular sides of hNTCP, respectively. The grey arrows are added according to the order of substrate accesses from extracellular side to intracellular side, thus constructing the substrate transport cycle of hNTCP.

### Coupling between Na^+^-binding and TCA-binding via X-motif conformational change

As hNTCP is a sodium-dependent secondary symporter, we investigated the role of Na^+^ in its substrate transport. In the three 5 μs MD simulations for TCA-bound hNTCP, two Na^+^ were built to coordinate with X-motif due to the reported importance of Na^+^ on the substrate transport^28, 46^. Structural analyses shows that all the stable TCA-bound conformational states are coupled with two Na^+^ binding to the negatively charged X-motif (Supplementary Fig. 10). Except for the key residues (Q68, S105, N106, T123, E257, and Q261) hypothesized by evolutional analysis, residues G101, G102, and S119 at the Na1-site, and S99 and C260 at the Na2-site also contribute to the stable binding of two sodium ions (Supplementary Fig. 10c). Time evolution of distances between Na^+^ and coordinated oxygen atoms reveals that these residues in the X-motif subtly cradle the two Na^+^ ions. We then carried out two 5 μs MD simulations each initialized with two Na^+^ but no TCA binding to the two experimentally determined conformations. Two Na^+^ ions were found to be stable during the 5 μs open-pore hNTCP simulation. However, the Na^+^-binding sites in inward-facing hNTCP became unfavorable without TCA as both Na^+^ escaped from their binding sites and the one initialized in the Na1-site entered the cytoplasm solvent after 500 ns (Supplementary Fig. 11).

In addition, we performed three 3 μs MD simulations each initialized with previously sampled stable TCA-binding conformations but the two Na^+^ removed. Without Na^+^-binding, the binding stability of TCA significantly decreased in all three simulations (Supplementary Fig. 12). In particular, TCA bound to the middle and bottom positions (binding poses 2 and 3) quickly underwent rotation after a few hundred nanoseconds, allowing its hydrophilic sulfate head group to access the water environment (Fig. 5a-c), which suggests a synergistic relationship between Na^+^-binding and substrate binding. In the absence of Na^+^ occupancy in hNTCP, the TCA substrate displayed common amphiphilic properties and oriented in the same direction as lipid molecules. Notably, the phenomenon of substrate flipping during transport has been observed in some transporters, most commonly cholesterol and fatty acids transporters. Recent studies have provided evidence of lipid flipping in a proton-dependent lipid transporter and an omega-3 fatty-acid transporter^47, 48^, suggesting that amphiphilic substrate may need to flip to maintain the same orientation relative to the surrounding local environment of the membrane bilayer.

**Fig 5.**
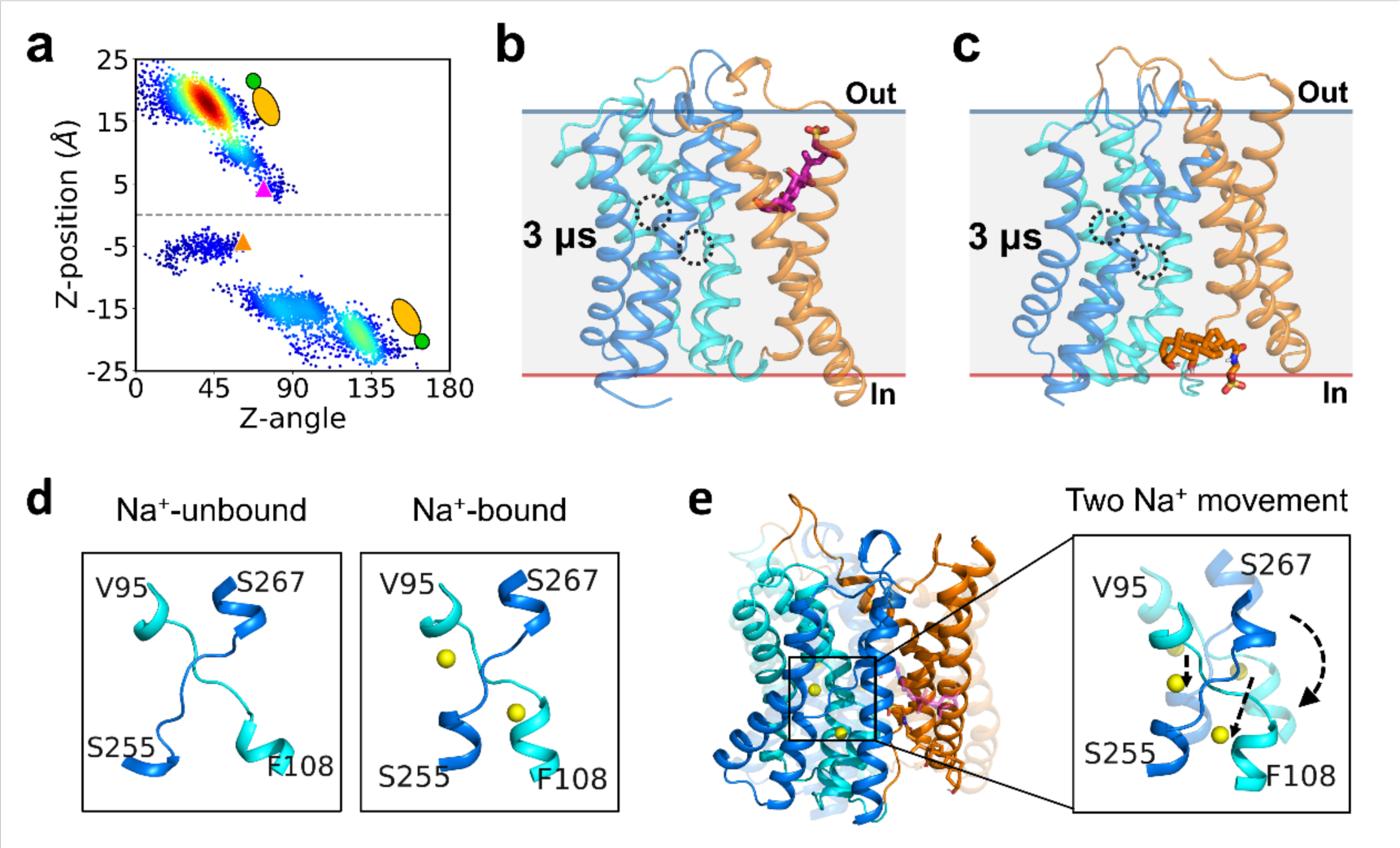
Coupled Na^+^-binding is essential to maintain the stable TCA-bound states in hNTCP. **a** Position and direction distribution of TCA in hNTCP without Na^+^-binding in the 3 μs MD simulations starting from binding pose 2 (magenta triangle) and 3 (orange triangle). **b-c** The poses of TCA in the extracellular pocket (**b**) and intracellular pocket (**c**) after 3 μs MD simulations without Na^+^-binding. The two dashed cycles highlight the two Na^+^-binding sites, in which no Na^+^ is placed. **d** Structural comparison of X-motif altered by Na^+^-binding. **e** The position shift of the two bound Na^+^ related to the conformational change of the folded X-motif between the open-pore and the inward-bound states.

It is interesting to note that there is little direct interaction between the two Na^+^ ions and TCA. Energy calculations along MD trajectories with both Na^+^ and TCA stably bound shows that the total interaction energy is less than −0.1 kcal mol^−1^ in all three binding poses (Supplementary Fig. 13). Structure clustering and comparisons reveals that the major effect of Na^+^-binding is the tightening of the X-motif (Fig. 5d), which is well-coordinated with the sulfate head of TCA. The X-motif assumes a tightly folded state with Na^+^-binding, whereas it is more unwound and flexible without Na^+^-binding, reducing its ability to form stable contacts with TCA. Further interaction energy calculations illustrate that Na^+^-binding indirectly enhances the interaction between hNTCP and TCA with energies changed by −40±6 kcal mol^−1^ upon Na^+^-binding and confirms that the changes are dominated by the interaction energies between the X-motif and TCA (Supplementary Fig. 14 and 15). Our results suggest that Na^+^-binding induces the conformational change of X-motif and thus modulates the local environment for substrate binding in hNTCP. The delicate cooperation among Na^+^-binding, X-motif conformational change, and TCA-binding in hNTCP is also implied in the alignment of the Na^+^-bound X-motif in the open-pore and the inward-facing hNTCP (Fig. 5e). Transition from the open-pore to the inward-bound conformation accompanies the movement of two Na^+^ ions further towards cytoplasm and minor conformational shifts of the folded X-motif.

### Na^+^-binding drives substrate translocation in hNTCP

After establishing that Na^+^-binding impacts substrate binding in hNTCP, we further investigate the effect of Na^+^-binding on the thermodynamic properties of substrate translocation with enhanced sampling MetaD simulations. Four sets of well-tempered 2D-MetaD simulations were initialized using the stable TCA-bound outward-facing and the inward-facing state, each with or without Na^+^-binding (Supplementary Table 1). The Z-position and Z-angle of TCA were used as CVs to sample its translation and rotation in hNTCP. The resulted free energy profiles of TCA translocation in the extracellular and the intracellular pockets were constructed under different Na^+^ binding states (Fig. 6). Convergence of these results was checked using two additional sets of independent replicates (Rep. 1 and Rep. 2) initialized with different equilibrated conformations and random seeds for initial velocities (Supplementary Fig. 16). Together with the Rep. 0 results shown in Fig. 6, the evident alteration in the shapes of the free energy profiles in all three independent replicates reveals the determining role of Na^+^-binding on consecutive TCA translocation.

**Fig 6.**
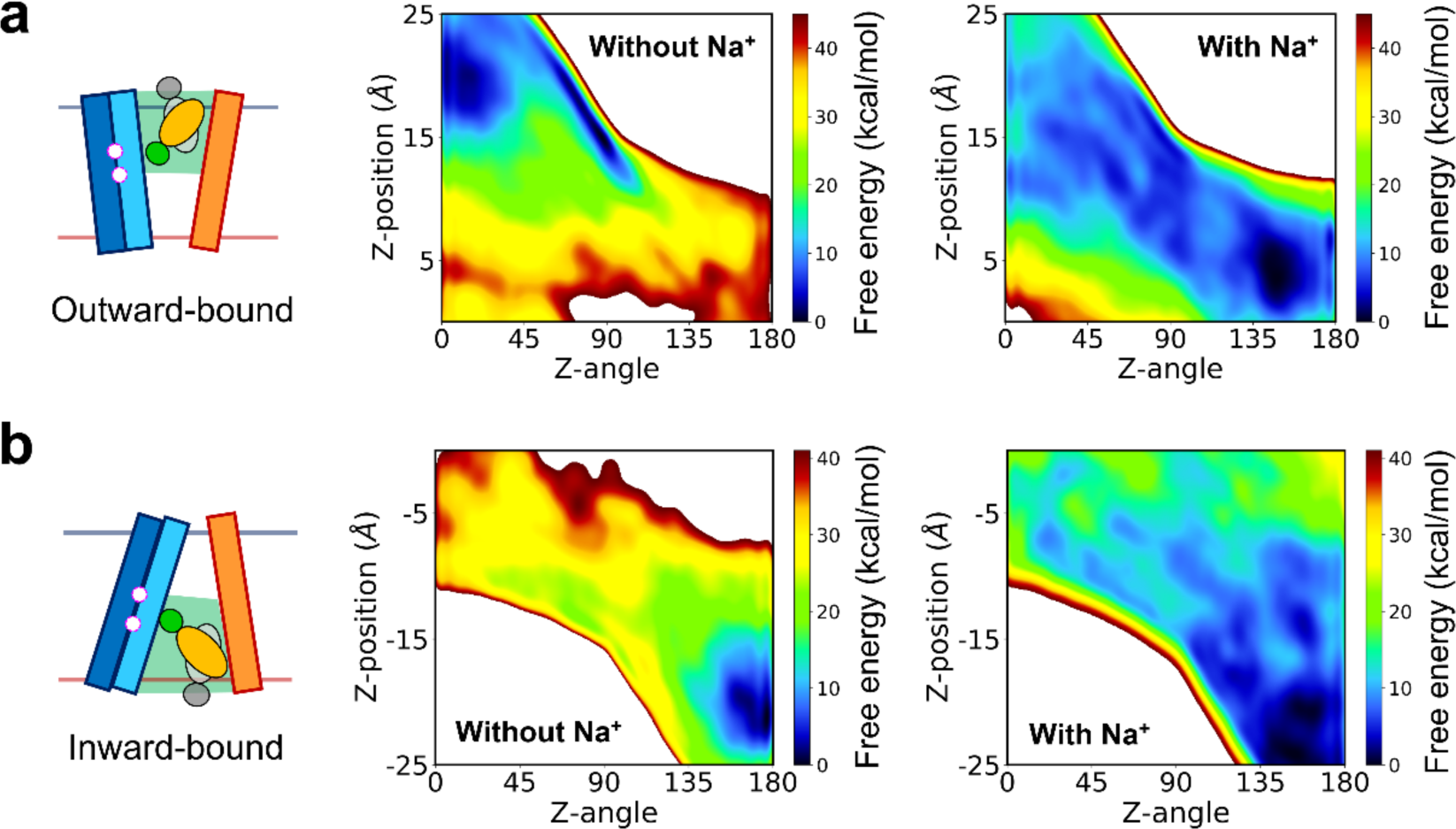
How Na^+^-binding drives the substrate transport. **a** The shape of the free energy surface of TCA translocation in the extracellular pocket is reversed by Na^+^-binding, with the hydrophilic sulfate head group shifting from the extracellular water environment and translocating into the middle pore. **b** The free energy surface of TCA translocation in the intracellular pocket is reshaped by Na^+^-unbinding, with fast downhill process of TCA release.

When TCA is located in the extracellular pocket, the shape of the free energy surface is completely reversed by Na^+^-binding, with the global minimum shifting from the hydrophilic sulfate head group being close to the extracellular water environment (Z-position=19.8 Å) to it translocating into the middle pore and coordinating with the X-motif (Z-position=3.6 Å) (Fig. 6a). Without Na^+^-binding, TCA is most favorable with a head-up, tail-down configuration (Z-angle around 0°), which is presumably how the TCA diffuses into the pocket. After Na^+^ ions enter their binding sites at the X-motif, a new free energy minimum emerges that is 10.2±1.1 kcal mol^−1^ more favorable, providing the thermodynamic incentive for TCA to orientate into a head-down, tail-up configuration with Z-angle around 145° (binding pose 1). Our simulations reveal that Na^+^-binding to X-motif activates TCA transport.

When TCA is in the intracellular pocket, the global minima in the free energy profiles are similar in both Na^+^-binding states, with the sulfate head of TCA accessing the intracellular water environment. This suggests that the inward-bound state leads to spontaneous release of the substrate. The difference in the free energy landscapes again highlights the importance of Na^+^-binding in this process (Fig. 6b). When the Na^+^-binding sites were occupied, the free energy landscape is relatively flat with multiple minimum and barriers in between, such that the transition would be a relatively slow process. When there is no Na^+^-binding, the configuration with Z-position=-5.6 Å, Z-angle=43° (binding pose 3) becomes significantly more positive in free energy than the global minima, leading to a fast downhill process for TCA dissociation. The results from MetaD simulations are consistent with the observations from microsecond conventional MD (cMD) simulations of systems 3 and 9 (Supplementary Table 1), both suggesting that the dissociation of Na^+^ facilitates the reorientation of TCA for its translocation in hNTCP.

Many biochemical and biophysical studies have indicated that sodium ions provide the driving force for the transport of various substrates such as glucose, amino acids, sulfate/carboxylic acid, and anions like Cl^−^/I^−50^. The mechanistic details have been investigated at the atomic level using computer simulations. For instance, research on the role of sodium ions in Glt_Ph_ illustrated that subtle structural rearrangements following the release of a sodium ion at site 2 induce the reorientation and translation of the substrate aspartate, thus allowing the coupled release of the sodium ion and the substrate into the cytoplasm^34^. A similar mechanism has been demonstrated in the computational study of LeuT, in which the coupled Na^+^- and substrate-binding events are accompanied by both local and global (interhelical) rearrangements in the outward-facing state of LeuT^35^. Our previous study on SLC26A9 indicated that Cl^−^ binding is coupled to the Na^+^-binding^51^, a finding also corroborated in studies of the cation-chloride cotransporters SLC12 family^52^. Recently, Han et al. combined all-atom MD simulations with structure studies to examine SGLT1, found out that the glucose coordination depends on the binding states of two sodium ions^37^. In particular, they observed that glucose exhibits greater mobility when there is no sodium in the binding pocket. Our study extends this body of knowledge by elucidating the coupling mechanism of bile acid and two Na^+^ in hNTCP. We clarify the coordination mode of Na^+^ and TCA in alternative conformational states of hNTCP through structural dynamic analysis, and also illustrates that Na^+^-binding drives substrate translocation via thermodynamic analysis.

### Substrate translocation relies on alternative-state transition to access the intracellular side

The transition between the Na^+^-binding, outward-bound state and the Na^+^-binding, inward-bound state were further studied using MetaD simulations. The relationship between local gating motions and global structural transitions is a crucial aspect of the transport mechanism, and two sets of well-tempered 2D-MetaD simulations were performed over 900 ns to shed light on this topic (Supplementary Table 1). The Z-angle of TCA were used as one CV to describe the substrate movement, while the conformations of hNTCP were characterized by the native contact differences (*δQ*) in TMD as another CV. In particular, the native contact difference *δQ*_OP-OF_ was used to characterize the conformations along the transition path from outward-facing to open-pore, and *δQ*_IF-OP_ was used to characterize the conformational transition from open-pore to inward-facing (Fig 7a, see Methods). The construction of free energy surface profiles reveals the relationship between TCA translocation and the conformational changes of hNTCP. We note that *δQ*_OP-OF_ and *δQ*_IF-OP_ are two distinct CVs for driving different conformational sampling, and the corresponding 2D projection of free energy surfaces are not comparable.

**Fig 7.**
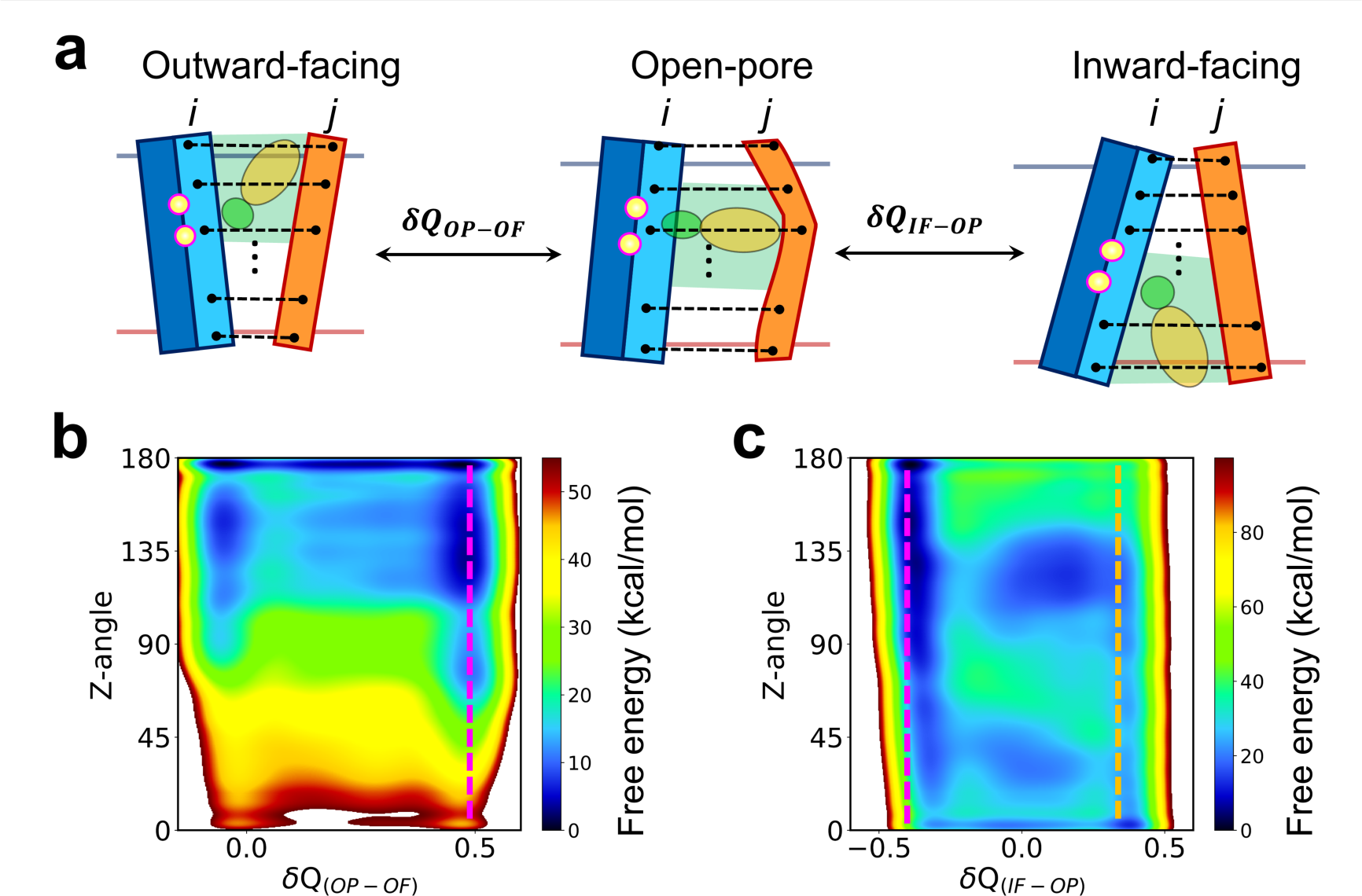
Substrate translocation relies on alternative-state transition to access the intracellular side. **a** Scheme of 2D-MetaD simulations for the conformational transition of the alternative states of hNTCP. **b** The free energy profile of outward-bound to open-pore transition (*δQ*_OP-OF_). **c** The free energy profile of open-pore to inward-bound transition (*δQ*_IF-OP_). The magenta and orange dashed lines mark the open-pore and the inward-facing cryo-EM structures, respectively.

The free energy profile of the outward-bound to open-pore transition shows that the outward-bound and open-pore states of hNTCP are both thermodynamically favorable (Fig. 7b). The transition from the outward-facing to the open-pore conformation does not need to involve the reorientation of TCA and is thermodynamically favored. When the hNTCP assumes the open-pore conformation, the TCA can switch between two orientations which has Z-angles of 135° and 85°, respectively. Based on the free energy profile of open-pore to inward-bound transition, a slight conformational shift towards the inward-facing conformation allows TCA to rotate further down with a Z-angle smaller than 90° (Fig. 7c). The open-pore state is thermodynamically more favorable than inward-bound state when the hydrophilic sulfate head group of TCA interact with X-motif, and an intermediate minima was observed at *δQ*_IF-OP_ ∼ 0. The transition entails more subtle coupling between the substrate orientation and protein dynamics and involves overcoming multiple energy barriers. Our free energy calculation results are consistent with the fact that the outward-facing conformation was not captured in structural biology experiments, while the open-pore and the inward-facing conformations were. Taking together, the global conformational transitions of hNTCP that enable TCA transport are examined from a thermodynamic perspective, facilitating to the construction of transport cycle.

### Overall scheme of the hNTCP transport cycle

Earlier, the elevator-type alternating-access model was presumed for several transporter families including SLC10^27, 53^. In this mechanism, substrate translocation accessibility is obtained by overcoming a fixed barrier of one domain movement against another relatively rigid domain, with substrate binding and release in each alternative state facilitated by local gating transitions in the moving domain. This elevator-type alternating-access mechanism is only partially relevant to hNTCP; the extensive MD simulations, starting with experimentally resolved open-pore and inward-facing structures, provide a more vivid picture on its transport cycle. Combining the dynamic and thermodynamic evidence in apo and TCA-bound hNTCP, a complete molecular mechanism of substrate transport is proposed (Supplementary Movie 1) including the following key steps:

1. Under Apo condition, hNTCP is intrinsically dynamic and populates a range of conformations including outward-facing and inward-facing ones, as evidenced in three 5 μs cMD simulations (Supplementary Table 1, MD systems 4-6). When apo-hNTCP adopts outward-facing configurations, amphiphilic molecules such as TCA can access into the extracellular pocket but cannot directly enter the middle pore of TMD for further transport.
2. Upon Na^+^-binding, substrate translocation is thermodynamically activated as TCA can flip from the extracellular side into the middle pore where it stably stays (MD systems 1-2, 7-8, 12, 14). Na^+^-binding enhances the coordination of the X-motif with the sulfate head group of TCA by tightening it up.
3. The Na^+^-bound and TCA-bound hNTCP undergoes conformational transition from the outward-facing to the open-pore conformation. This open-pore state is stable and provides space for the rotation of the TCA tail group to occur (MD systems 12, 14, 16-17).
4. The hNTCP transitions into an inward-facing conformational state as the sterol tail of TCA flips down. With Na^+^ bound, TCA maintains stable in the inward-facing hNTCP on the microsecond timescale (MD system 3).
5. Finally, Na^+^-unbinding facilitates the detachment of TCA from the intracellular pocket of hNTCP. With the release of the two Na^+^ ions into the cytoplasm, TCA flips again, exposing its sulfate head group to the cytoplasmic solvent (MD systems 13-15). The release of TCA allows hNTCP to return to the dynamic-apo state in step (1) and restarts the transport cycle.

## Discussions

The hNTCP has been established as a valuable target for combatting HBV/HDV infection^14^, as one of its macromolecule inhibitors, myrcludex-B, entered clinical trials^54^. In parallel, many efforts have been made to develop small molecule NTCP inhibitors through virtual screening, high-throughput screening, and drug repurposing ^55–58^. The design of these inhibitors primarily focuses on disrupting the recognition of HBV by hNTCP and has limited consideration for preserving the intrinsic transport capacity, probably due to the lack of structural and mechanistic understandings of hNTCP. Current hNTCP inhibitor design strategies include jamming the recognition interface on the extracellular side of the core domain or competing with the first 48 residues of myr-preS1 for binding at the extracellular pocket. Either may seriously impact the intrinsic transport function of hNTCP *in vivo*. The timely insights into the transport mechanism by this work are expected to accelerate the development of hNTCP-targeted therapies.

The hNTCP is generally considered as an elevator-like transporter, based on its conformational symmetry and rigidity^15, 27^. Over the years, the inward-facing conformation of elevator-like transporters has been most frequently determined, suggesting that the probabilities of other states are relatively low in the conformational ensemble^45^. The successful cryo-EM determination of the open-pore and inward-facing hNTCP structures was achieved through the use of Fab fragments. Conversely, without the stabilization effect of antibody binding, apo hNTCP is highly dynamic, as demonstrated by large-scale MD simulations. Meanwhile, through molecules docking and MD simulations, multiple binding events have been identified corresponding to the outward-facing, open-pore, and inward-facing states of hNTCP.

Our results further indicate that in its apo state, the middle-pore of hNTCP is closed, and the experimentally resolved open-pore structure actually represents a substrate-bound conformational state. Interestingly, open-pore conformation is very uncommon as an intermediate state in the transport processes of known transporters, as most transporters seldom exhibit significant changes in the backbone of the central region of their transmembrane helices^27^. For small substrate molecules, the intermediate state in which the substrate passes halfway through the transport pathway can be achieved by altering the local conformation of several key residues, as observed in transporters such as SGLT, LeuT, KCC and NKCC^30–38^. For relatively large substrate molecules such as TCA, the transmembrane helices need to be rearranged to form an open-pore state to provide sufficient space for the substrate to pass through, as observed also in lipid transporters^47, 48^.

One of the most intriguing questions for sodium-dependent secondary transporters is how the sodium ions drive the transport processes. Na^+^ can play diverse roles in membrane transport. For example, the importance of the sodium ion was found to be structural for maintaining the homodimer assembly of Fluc^46, 71^, a sodium-dependent fluoride channel, although a recent study challenged this idea and suggested that Na^+^ binding directly activates the F^−^ transport^72^. In the case of LeuT-fold transporters, sodium ions at two distinct binding sites are thought to play different roles, with one Na^+^ directly coordinating with the substrate and the other primarily coordinating with backbone and side chain atoms on the TM helices to regulate gating^32^. All-atom MD and MetaD simulations offer the opportunity to examine the dynamic and thermodynamic impact of sodium ions with *in silico* experiments by varying simulation conditions. We uncover a unique mechanism of Na^+^ binding that facilitates transport by flipping the substrate, exploiting its amphiphilic nature. We also show that this effect is exerted indirectly by coupling with the conformation change of the X-motif in hNTCP and modulating the binding environment for the substrate.

As a secondary transporter, hNTCP utilizes the sodium electrochemical gradient to transport substrates across the plasma membrane. In this study, we assume that the sodium ions flow from the extracellular side (high concentration) to the intracellular side (low concentration) and didn’t explore the atomistic details of sodium entry and release. As there is 150 mM NaCl homogeneous distributed in the simulation systems, we examined potential Na^+^ entry events or the presence of extracellular Na^+^ high-frequency access sites in the three 5 μs apo-hNTCP MD simulations. However, no Na^+^ entry was observed and no significant high-frequency access site in hNTCP was identified in our simulations. We hypothesize Na^+^ passively diffuses from high concentration to low concentration following the water pathway as it should be hydrated during entry. A continuous hydration pathway was identified at the interface between the core and the panel domains, which connects the sodium binding sites at the X-motif and the extracellular solvent (Supplementary Fig. 17a). Two high-density hydration sites were identified along this pathway, of which the surrounding residues might be relevant to the Na^+^-selectivity of hNTCP (Supplementary Fig. 17b). Further biochemical and biophysical investigation is needed to fully understand this aspect of hNTCP transport.

We note that our simulations were based on monomic hNTCP, and the impact of oligomerization on its functional activity remains unclear^73, 74^. Previous studies have shown that oligomerization is essential during the maturation of hNTCP, and it occurs at the endoplasmic reticulum membrane of the secretory pathway and persists at the plasma membrane. However, the effect of oligomerization on the intrinsic function of hNTCP is yet to be studied. Our simulations suggest that monomic hNTCP can carry out the entire substrate translocation cycle. A recent work proposed a dimeric model of NTCP with the dimer interface consisting of TM4, TM7, TM9 and the full C-terminus^75^. This interface is located far away from the extracellular and intracellular pockets, implying that oligomerization may not significantly impact substrate transport.

Interestingly, the alignment of the predicted dimer interface with a cryo-EM monomer structure (PDB id: 7ZYI)^18^ reveals electron densities that likely correspond to substrates at the interface (Supplementary Fig. 18). This observation suggests that oligomerization may play a role in substrate recruitment. Indeed, how bile acids are recruited by hNTCP despite their low extracellular concentrations remains elusive. A plausible hypothesis is that the amphiphilic substrate diffuses in the outer leaflet, while hNTCP potentially fosters a favorable local environment for substrate recruitment and access to the extracellular pocket. Whether sodium binding facilitates this process is also uncertain and further investigation will be needed.

In conclusion, this work represents a significant step towards uncovering the molecular mechanisms of hNTCP at the atomistic level by constructing its transport cycle and illustrating how the sodium ions modulate the free energy landscape of substrate translocation. The complexity of hNTCP, including endogenous substrate transport pathway, sodium pathway, recognition site for HBV, and dimerization, must all be considered when developing potent and safe hNTCP-targeted therapies. It would be a daunting task to design therapeutics that minimize negative side effects by preserving the intrinsic function of bile acid uptake. Advancing our understanding of the molecular mechanisms of hNTCP is crucial for successfully overcoming this challenge.

## Conclusion

MD simulations, cumulatively more than 50 μs, were employed to study the conformational dynamics of hNTCP during TCA transport. To gain thermodynamic insights into the conformational changes, 2D-MetaD simulations were used to construct free energy profiles among different states. Our computational approach allows us to demonstrate that the experimental open-pore structure is a substrate-bound structure and elucidate the binding mode. We found that Na^+^-binding activates the TCA translocation by flipping it from the extracellular side to the middle pore where it remains stably anchored. This is accomplished by tightening the X-motif to better coordinate with the sulfate head of TCA. Na^+^-release promotes rapid TCA detachment from inward-facing hNTCP. Free energy profiles were constructed along the TCA translocation pathway, highlighting the role of Na^+^ in modulating the protein electrostatic environment for substrate reorientation. Our results show that TCA translocation relies on the conformational transition of hNTCP from the outward-bound state to the most stable open-pore state and to the inward-bound state, while the inward-bound state is flexible with a greatly deformed intracellular gate and a less stable TCA. Structural analysis reveals that the conformational rearrangements of hNTCP in the transport cycle mainly occur at the interface between the core and the panel domains, involving TM1, TM3b, TM5, TM6, TM8a and TM9. Taken together, we uncover an interesting mechanism of Na^+^-driven transport in hNTCP that leverages the amphiphilic nature of the substrate.

## Methods

### Preparation of substrate-bound wild type hNTCP structures

To investigate the intrinsic transport mechanism of hNTCP, we prepared multiple models for apo and holo hNTCP structures. Two representative open-pore (PDB id: 7PQQ) and inward-facing (PDB id: 7PQG) conformations of hNTCP were used to set up the simulation systems. The mutated residues in the cryo-EM structures were restored to the wild type, including mutations F33V, F37I, K86N, V95I, V107I, C129L, and L221H. The binding modes of the two crucial Na^+^ ions were manually built based on structural analyses and then optimized by energy minimization to equilibrate the local environment. Protonation states of polar residues were calculated with PropKa^76^.

Complex structures corresponding to the open-pore and inward-facing conformations were generated using molecular docking with the MOE package^77^. Docking pockets were defined for these two conformations by all residues within 5 Å of N103, Q264, and S199. The three-dimensional structure of substrate TCA was generated and subjected to energy minimization using MOE. For induced-fit docking in each conformation, one hundred ligand conformations were generated using the ’London dG’ scoring function. These were subsequently refined by molecular mechanics and rescored with the ’GBVI/WSA dG’ scoring function to identify the top five docking poses with the highest docking scores^77^. Finally, alignment and clustering of the top five docking poses yielded two dominant docking poses for the open-pore hNTCP (docking poses 1 and 2) and one docking pose for the inward-facing hNTCP (docking pose 3). To investigate the effect of Na^+^-binding, the stable complex structures after 5 μs MD simulations were collected to alter the Na^+^-binding states and perform MD simulations for Na^+^-unbound complexes. A total of eleven structures were prepared to set up MD simulations (Supplementary Table. 1, MD systems 1-11).

### MD simulation systems setup and equilibration

Using the CHARMM-GUI protocol for building bilayer membrane system^78, 79^, we embedded the prepared structures into a 1-palmitoyl-2-oleoyl-sn-glycero-3-phosphocholine (POPC) lipid bilayer, with a total of pre-equilibrated ∼160 POPC molecules per simulation system. Then, we used TIP3P water molecules for solvation and 0.15 M NaCl for neutralization^80^. The position of hNTCP TMD in the lipid bilayer was obtained by aligning it with the structure 4N7W downloaded from OPM database^81^. For each system, the simulation box is approximately 82 x 82 x 100 Å^3^, containing a total of around 61,000 atoms. We note the exclusion of cholesterol molecules from the simulated membranes. Although cholesterol is physiologically present in the human membrane environment, it has been shown to bias the conformations of membrane peoteins^82, 83^. As such, the cholesterol molecules were excluded to prevent unintended inhibition of transporter dynamics^84^.

The CHARMM36m force field was used to describe the proteins^85^, and the parameters for substrate TCA were generated using the CHARMM general force field (CGenFF)^86^. The simulation systems were first relaxed through steepest-descent energy minimization. Solvent molecules in each system were relaxed via a 500 ps NPT simulation with restraints on all heavy atoms of solute molecules. Then 1 ns equilibration was established to relax the pre-equilibrated lipid bilayer, refining the conformations of lipid molecules around hNTCP. Subsequently, 2 ns equilibration was performed with gradually reduced restraint force on the heavy atoms of hNTCP, TCA, and sodium ions. Finally, 1 ns equilibration was carried out with force constant reduced to 0 kcal mol^−1^ Å^−2^ for all atoms in the MD simulation systems. All equilibrations were performed at a temperature of 310 K and a pressure of 1 bar.

### Conventional MD and 2D-MetaD simulations

All production simulations were performed using OpenMM package^87^ patched with the PLUMED 2.5 plugin^88^. Particle–mesh Ewald summation was used to calculate the long-range electrostatic interactions with a 12 Å real-space cut-off and periodic boundary conditions^89^. The Lennard–Jones interactions were truncated at 12 Å with an atom-based force switching function starting at 10 Å. The bonds involving hydrogen atoms were constrained using the SHAKE algorithm^90^. A time step of 2 fs was used in all simulations. Snapshots were saved every 20 ps for the protein and 1 ns for the entire simulation system via calling the MDTraj package^91^. In the conventional MD simulations, three independent replicates were performed to estimate the consistence of the dynamic properties. The duration of conventional MD simulations and 2D-MetaD simulations are provided in Supplementary Table 1.

The well-tempered approach was employed in all 2D-MetaD simulations^40^. For well-tempered MetaD, a bias factor of 0 is equivalent to standard MD a bias factor approaching infinity corresponds to standard metadynamics^40^. In between these extremes, we regulate the extent of conformational exploration by tuning the bias factor to avoid overfilling and to save computational time. The Gaussian height was set to 5 kJ mol^−1^, and the bias potential was added every 10 ps (5,000 MD steps), a time interval large enough for the conformation to relax^40^. This means that a Gaussian deposition rate of 0.5 kJ mol^−1^ ps^−1^ was initially used and gradually decreased. According to the corresponding fluctuations in the unbiased MD simulation, the Gaussian widths were set to 0.1, 4, and 0.1, respectively. Specific CVs used for 2D-MetaD simulations were defined using the MATHEVAL function in PLUMED and parameter details in MetaD simulations were summarized in Supplementary Table 2 and 3.

The choice of CVs is critical in metadynamics and must be selected carefully to ensure that the system explores the relevant phase space^92^. Appropriate CVs are expected to provide good separation between different states and substantially enhanced sampling^92^. To characterize the TCA bound state, the relative position of its sulfate head group along the membrane normal (Z-position) with the bilayer midplane at Z = 0 Å, and the angle between its tail-to-head vector and the normal vector of the membrane plane (Z-angle) were used to sample the TCA-bound conformational states (Fig. 2a). These CVs can well describe the TCA conformations and distinguish the different binding states of the substrate in the complex (Supplementary Fig. 2a), thus they were also used to drive the substrate movement in metadynamics (Supplementary Table. 1, MD systems 12-19).

To characterize the conformation transition states of hNTCP, the sizes of the extracellular gate, the middle pore, and the intracellular gate were used to sample the conformations of hNTCP, defined using selected residues listed in Supplementary Table 2. These three CVs effectively distinguish the different states of hNTCP (Supplementary Fig. 5a). To perform the efficient metadynamics with as few dimensions as possible, the conformational difference between the states was reduced to one dimension to drive the conformational transition of hNTCP. First, the native contact *Q*(*X*) of a conformation *X* was defined accordingly^93^

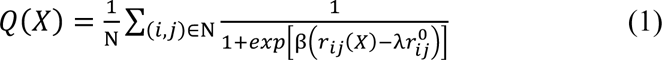

where N is the number of contact pairs of the target structure and 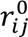 is the equilibrium distance between atom *i* and atom *j* in the unbiased MD simulation. β is a smoothing parameter with a value of 5 Å^−1^ and λ is set to 1.2.

Then, we selected a set of pairwise heavy atoms *ij* capable of distinguishing the three alternative conformations of hNTCP. Pairs with a standard deviation of 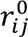 Larger than 3 Å among three conformations were selected, resulting in a total of 58 residues pairs (Supplementary Table 2). Subsequently, the native contact *Q*(*X*) of conformation *X* with reference to outward-facing state (*Q*_OF_ (*X*)), open-pore state (*Q*_OP_ (*X*)), and inward-facing state (*Q*_IF_ (*X*)) were calculated. The conformational transition of hNTCP along the transition paths of outward-bound to open-pore state and open-pore to inward-bound state were defined using native contact deviation *δQ*(*X*), written as

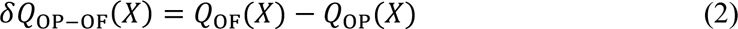

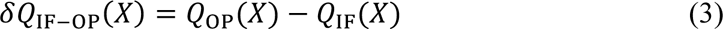

where *δQ*_OP-OF_ (*X*) represents the conformational transition of conformation *X* along the transition path from outward-bound to open-pore state, and *δQ*_IF-OP_ (*X*) represents the conformational transition of conformation *X* along the transition path from open-pore to inward-bound state.

### Simulation Analysis

VMD^94^ and MDAnalysis^95^ were used for the analysis of MD and 2D-MetaD trajectories, including geometry calculations, root-mean-square deviations (RMSDs), ligand-receptor interaction, etc. Hydrogen bonds were defined with a donor-acceptor distance cut-off of 3.2 Å and a donor-hydrogen-acceptor angle cut-off of 150°. The METAGUI 3 plugin^96^ was used to analyze the MetaD simulations. The free energy profiles were constructed by PLUMED, with the number of bins for the grid set to 500 to sum up the hills. The “mintozero” option was used to rescale the minimum value in bias to zero. Conformations in the local minima were extracted and clustered by METAGUI 3. All structural figures were generated using PyMOL (http://pymol.org/). Partial cryo-EM density in hNTCP was rendered by Chimera (https://www.cgl.ucsf.edu/chimera/). Continuous hydration pathway in hNTCP was rendered using iso-surface of water density in VMD.

To assess the quality of sampling in the well-tempered MetaD, we examined the time evolution of CVs to confirm that they propagated across relevant ranges in 2D-MetaD simulations. For each MetaD run, convergence was checked by analyzing the free-energy profile as a function of simulation time, thus determining whether the simulation time was sufficient^40^. In addition, we performed two extra independent replicates to construct the free energy profiles of TCA translocation in hNTCP. These sets of replicates were initialized with different equilibrated conformations and random seeds for initial velocities. Both strategies ensure that MetaD proceeds to convergence and the free energy results are qualitatively robust.

### Statistics and Reproducibility

Each simulation used the different random seeds to generate the initial velocity. No data was excluded from the sampled conformations and no blinding methods were used in simulation analyses. Statistical analysis was performed using NumPy.

## Data availability

The main data supporting the findings of this study are available within the article and its Supplementary Martial. The original data of MD simulations and generated force field parameters for TCA are available under a Creative Commons Attribution Non-Commercial 4.0 International license from https://doi.org/10.5281/zenodo.8220354. This defaults to the most recent version of the data.

## Code availability

All software used for this study are available and the links are provided as follows. For simulation systems setup: CHARMM c42b1 (https://www.charmm.org/archive/charmm/program/). For MD simulations: OpenMM 7.5.0 (https://simtk.org/projects/openmm), PLUMED 2.5 plugin (https://github.com/plumed/plumed2). For data analysis: VMD (http://www.ks.uiuc.edu/Research/vmd/), PyMOL (https://pymol.org/2/), METAGUI3 (https://github.com/metagui/metagui3), PLUMED-GUI (https://github.com/giorginolab/vmd_plumed), MDAnalysis package (https://www.mdanalysis.org/).

## Acknowledgments

The authors thank Prof. Jian Yang, Dr. Jing Xue, and Mr. Shufeng Sun for helpful discussions. The work is supported by the National Natural Science Foundation of China (32171247, 21803057), the “Pioneer” and “Leading Goose” R&D Program of Zhejiang (2023C03109), and the Westlake Education Foundation. We thank the Westlake University Supercomputer Center for computational resources and related assistance.

## Author Contributions

J.H. designed the experiment, analyzed the data, and wrote the paper. X.L. designed the experiment, performed the simulation, analyzed the data, and wrote the paper.

