## Supporting Information for "Molecular Mechanisms of Na^+^-driven Bile Acid Transport in Human NTCP"

Xiaoli Lu [1,2] and Jing Huang [1,2 *]

[1] Westlake AI Therapeutics Lab, Westlake Laboratory of Life Sciences and Biomedicine, 18 Shilongshan Road, Hangzhou 310024, Zhejiang, China.

[2] Key Laboratory of Structural Biology of Zhejiang Province, School of Life Sciences, Westlake University, 18 Shilongshan Road, Hangzhou 310024, Zhejiang, China.

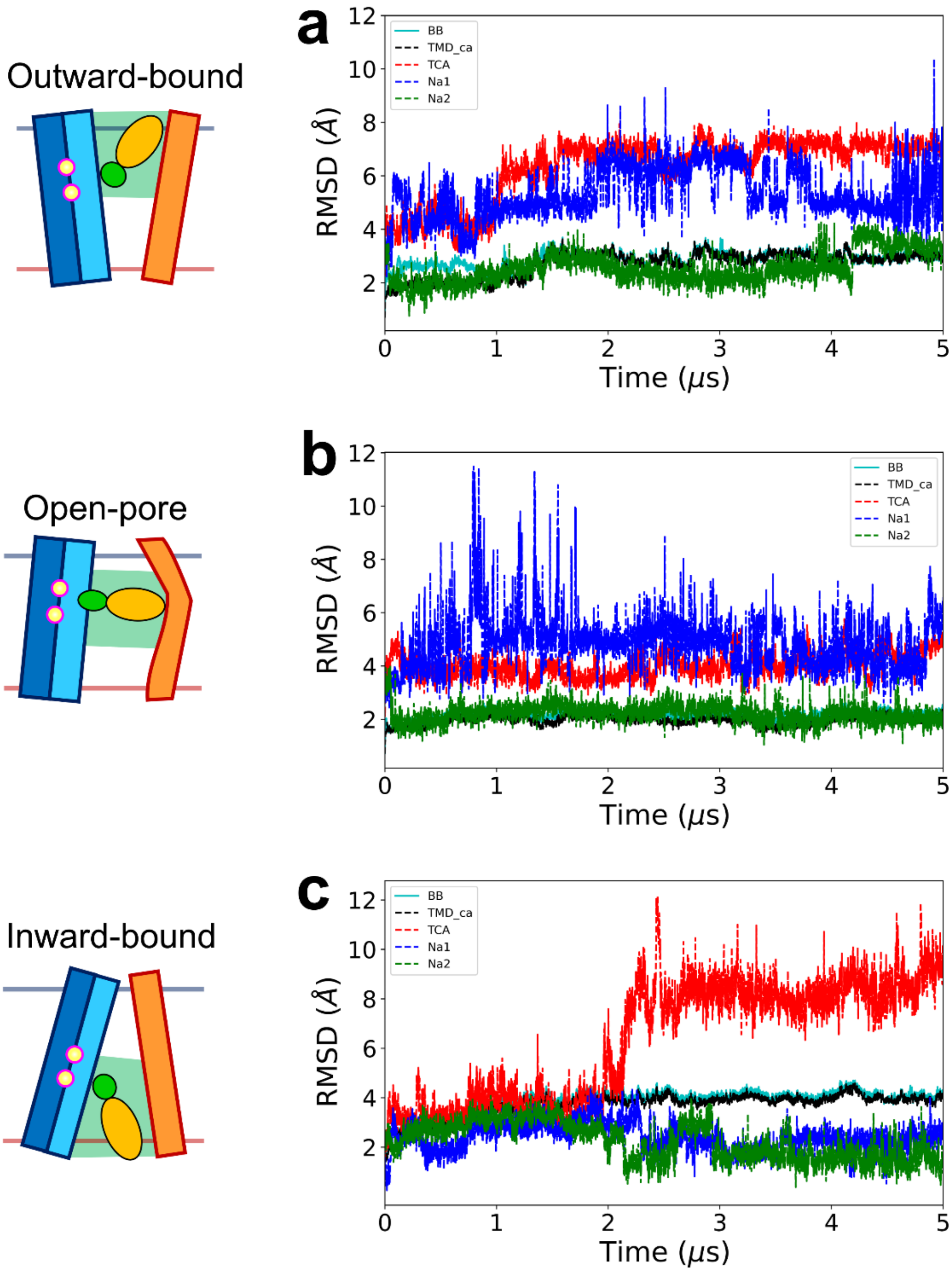

**Supplementary Fig. 1. Time evolution of RMSDs for TCA-bound hNTCP systems in 5 μs MD simulations. a-c** RMSDs of hNTCP TMD, substrate TCA, and two $Na^+$ in three TCA-bound hNTCP systems. The initial conformations of TCA-bound hNTCP are represented in the left cartoon.

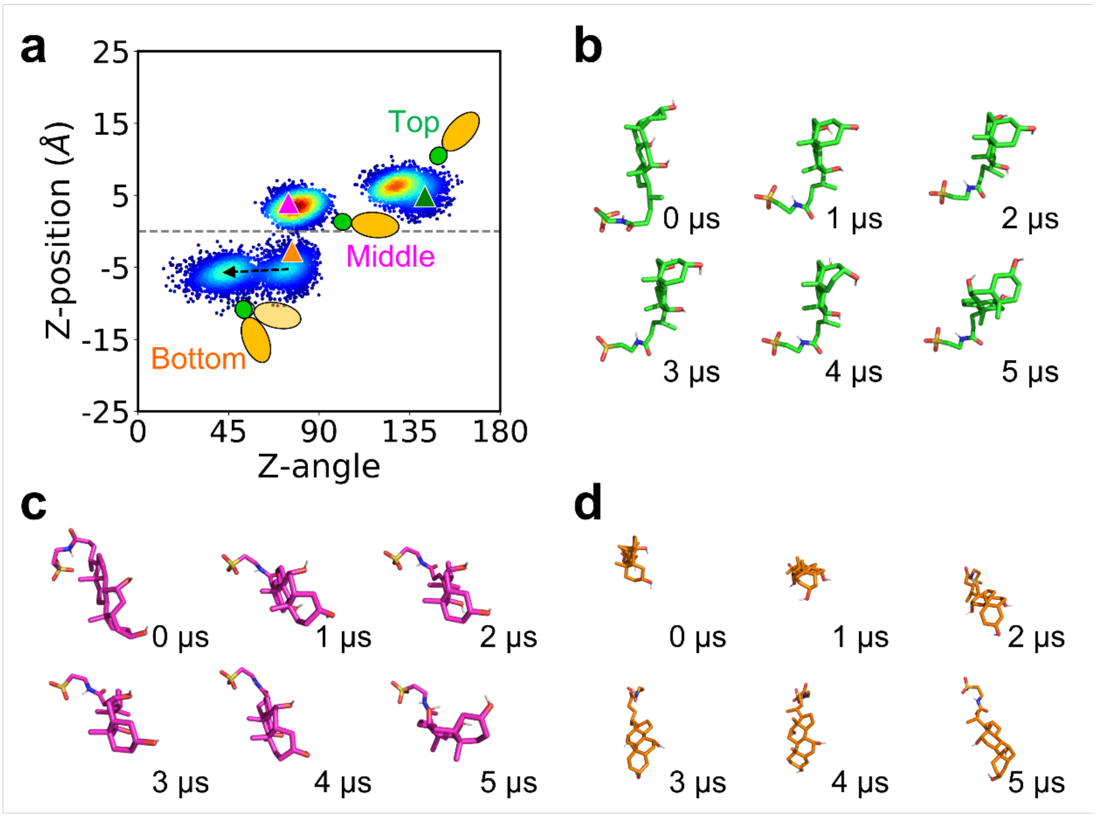

**Supplementary Fig. 2. The conformations of substrate TCA in hNTCP. a** Position and direction distribution of TCA in hNTCP with $Na^+$-binding from the three 5 μs MD simulations. The initial TCA-bound states corresponding to three docking poses 1-3 are marked with green, magenta, and orange triangles, respectively. The conformational fluctuations for the three states of TCA in the top extracellular pocket (green), middle pore (magenta), and the bottom cytoplasmic pocket (orange) are plotted together. **b-d** Snapshots collected every 1 μs to illustrate the transition of TCA poses to the stable binding poses in hNTCP. Alignment was performed with respect to hNTCP while the protein is hidden for clarity.

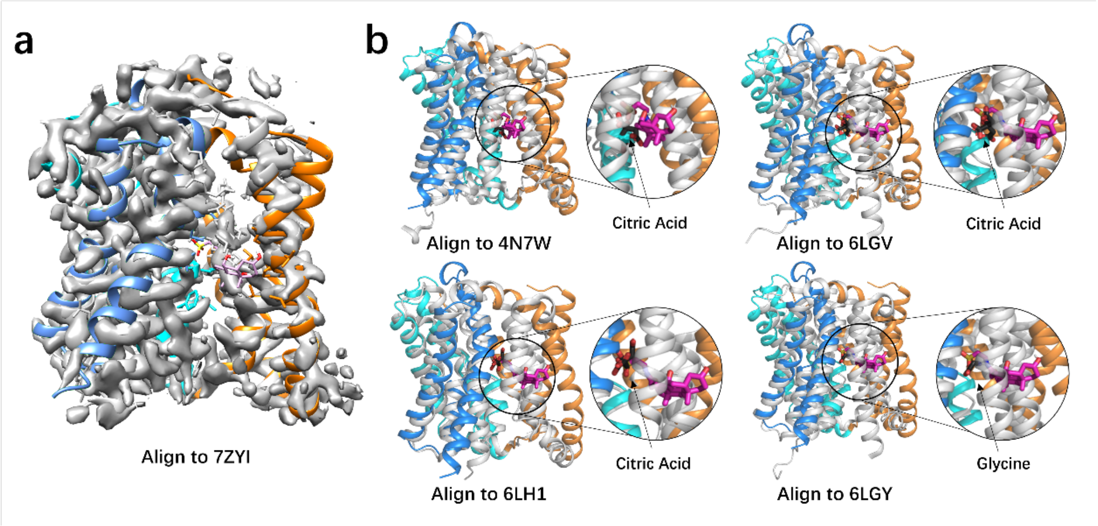

**Supplementary Fig. 3. Alignment of TCA-bound open-pore hNTCP with multiple experimental data. a** Alignment of TCA-bound open-pore hNTCP with the density of the complex structure of two GCDC molecules bound to hNTCP[1]. **b** Alignment of TCA-bound open-pore hNTCP with the structures of the citric acid or glycine (in black) bind to ASBT[2, 3].

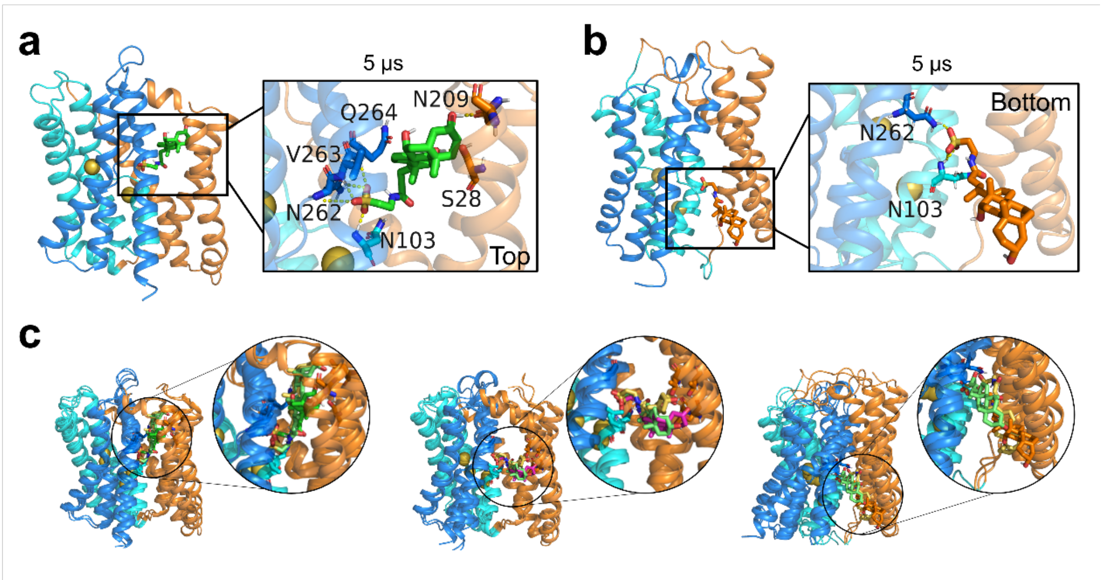

**Supplementary Fig. 4. Alternative binding modes between TCA and hNTCP captured from the 5 μs MD simulations with three replicates. a** The binding mode of TCA at the top position, where the lipophilic sterol tail group close to the extracellular side. **b** The binding mode of TCA at the bottom position, where the lipophilic sterol tail group close to the intracellular side. c. Alignment of each binding mode to the other two replicates 5 μs MD simulations. The binding modes in the extracellular pocket and middle pore are consistent in the position and involved residues. In intracellular pocket, binding modes from the different simulations are consistent in involved residues in X-motif but show differences in the position, which also indicates that binding in the intracellular pocket is unstable.

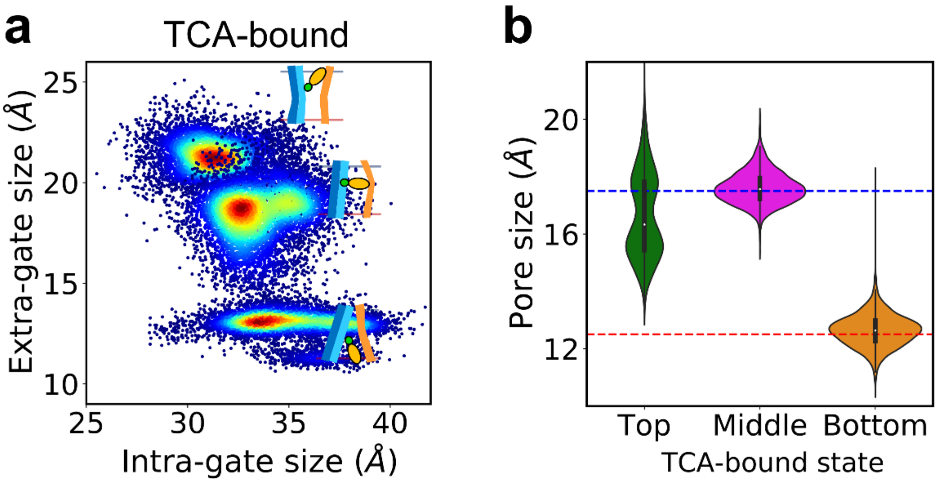

**Supplementary Fig. 5. The global conformational changes in the three 5 μs MD simulations for the three TCA-bound hNTCP systems. a** The size distribution of extracellular gate and intracellular gate of three TCA-bound hNTCP from the three 5 μs MD simulations. **b** The size distribution of the middle pore of three TCA-bound hNTCP from the three 5 μs MD simulations.

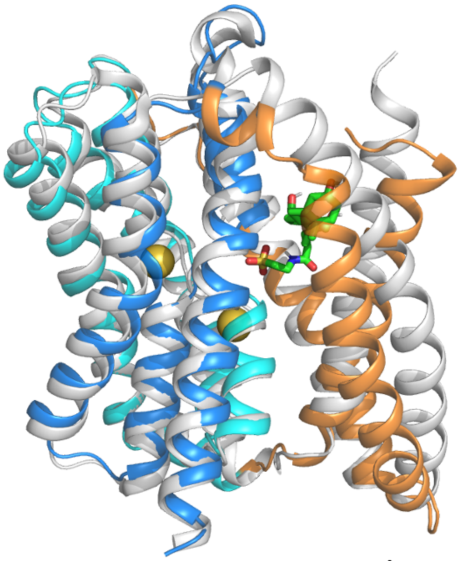

**Supplementary Fig. 6. Alignment of the outward-facing state sampled from MD simulations to the experimentally proposed outward-facing state.** The RMSD between two structures is 1.4 Å.

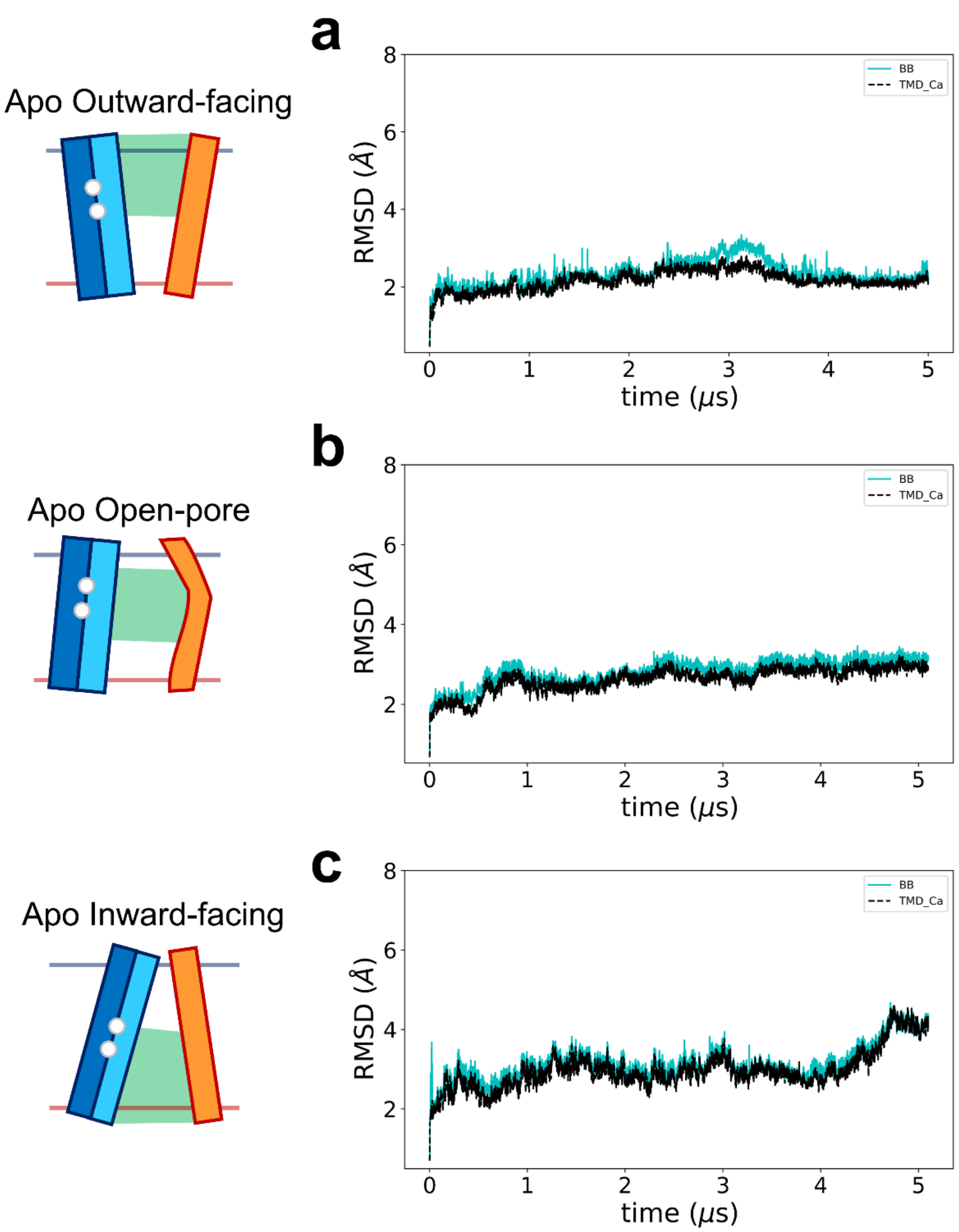

**Supplementary Fig. 7. Time evolution of RMSDs for apo hNTCP systems in 5 μs MD simulations.** The initial conformations of apo hNTCP are represented in the left cartoon.

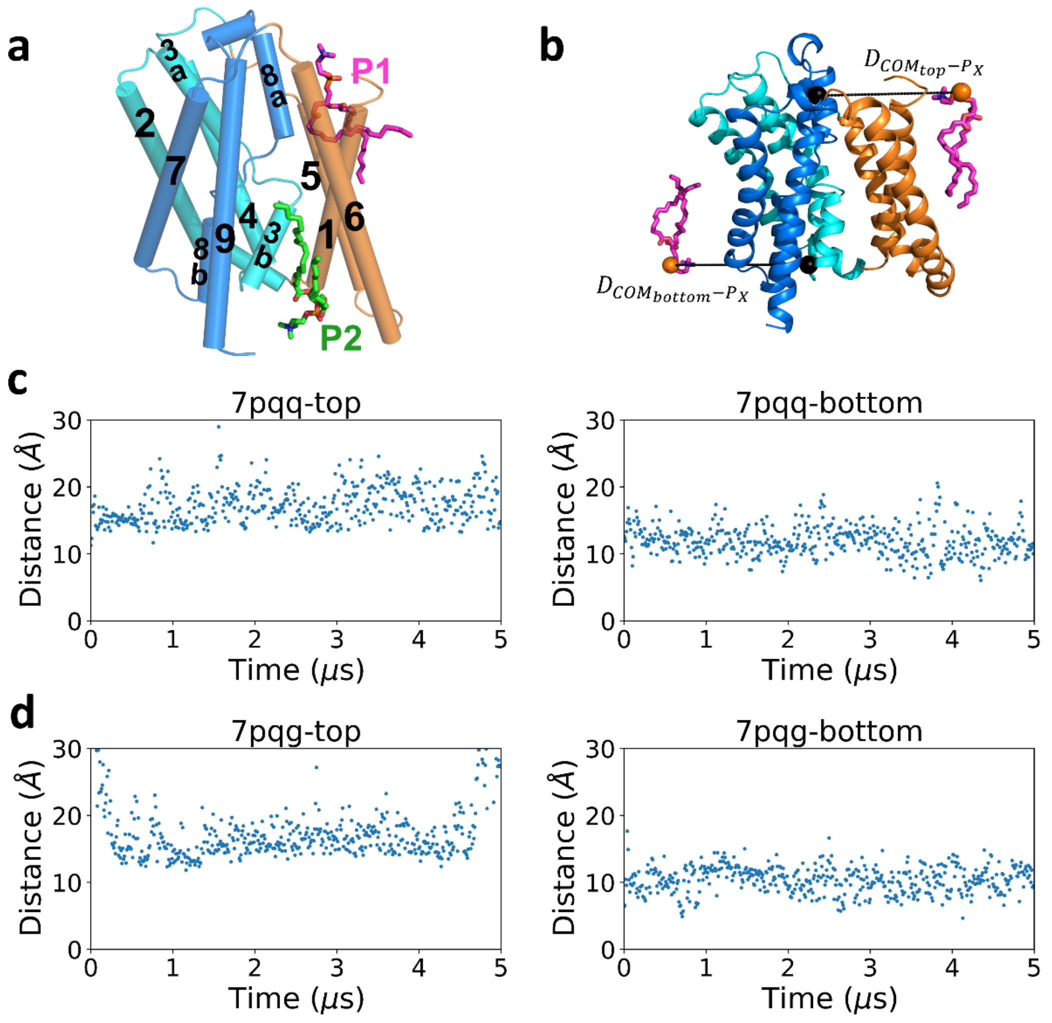

**Supplementary Fig. 8. Amphiphilic POPC can access the extracellular and intracellular pockets in 5 μs MD simulations for apo hNTCP systems. a** Two POPC molecules access the extracellular and intracellular pockets, respectively. **b** Schematic illustration of distance measurement to identify POPC molecules that interact stably with apo hNTCP. **c-d** Time evolution of distances defined in **b**.

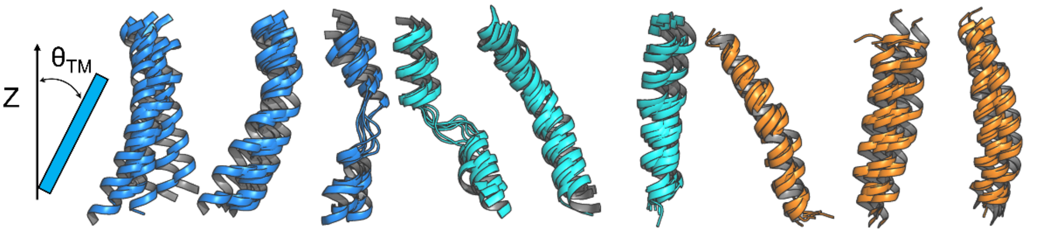

| | TM9 | TM7 | TM8 | TM3 | TM4 | TM2 | TM5 | TM1 | TM6 |
|---|---|---|---|---|---|---|---|---|---|
| $\theta_{TM}$ | 13±15 | 38±5 | 17±4 | 35±7 | 29±4 | 45±6 | 28±5 | 24±9 | 36±15 |

**Supplementary Fig. 9. The angle along the Z direction of nine TM helices of apo hNTCP in 5 μs MD simulations**.

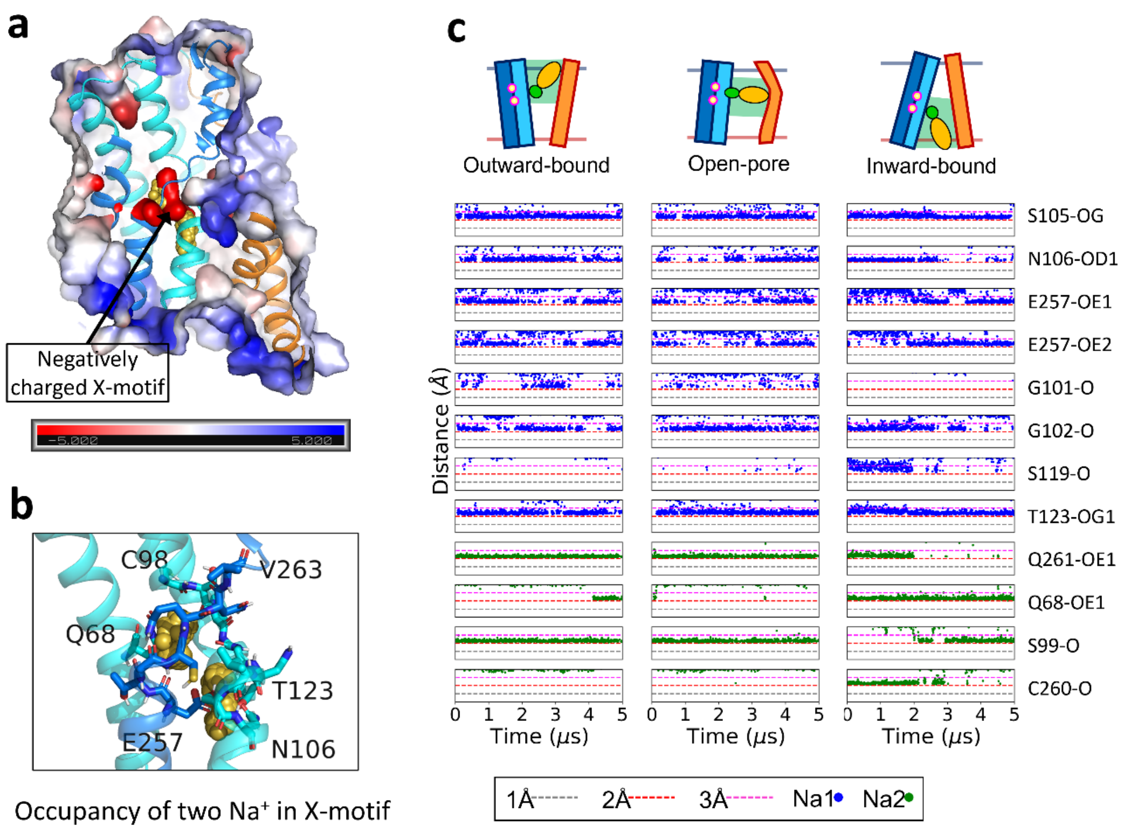

**Supplementary Fig. 10. Two Na$^+$ bind to a strongly negatively charged region. a** Surface representation of the electrostatic potential of hNTCP. **b** Na$^+$-binding occupancy in the X-motif formed by unwound regions of TM3 (C98 to N106) and TM8 (E257 to V263), cooperating with Q68 of TM2 and T123 of TM4. **c** Time evolution of distances between the two Na$^+$ and the residues involved in the coordination.

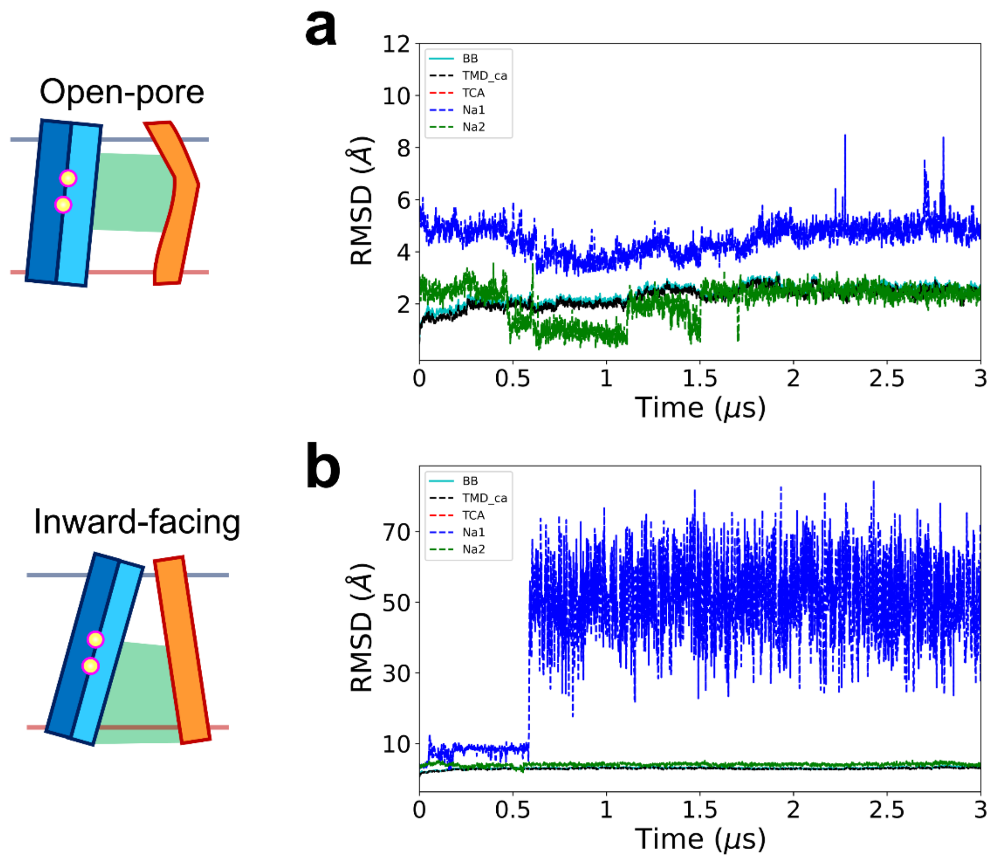

**Supplementary Fig. 11. Time evolution of RMSDs for $Na^{+}$-binding apo hNTCP systems in 3 μs MD simulations.** The two initial conformations of open-pore and inward-facing hNTCP are represented in the left cartoon.

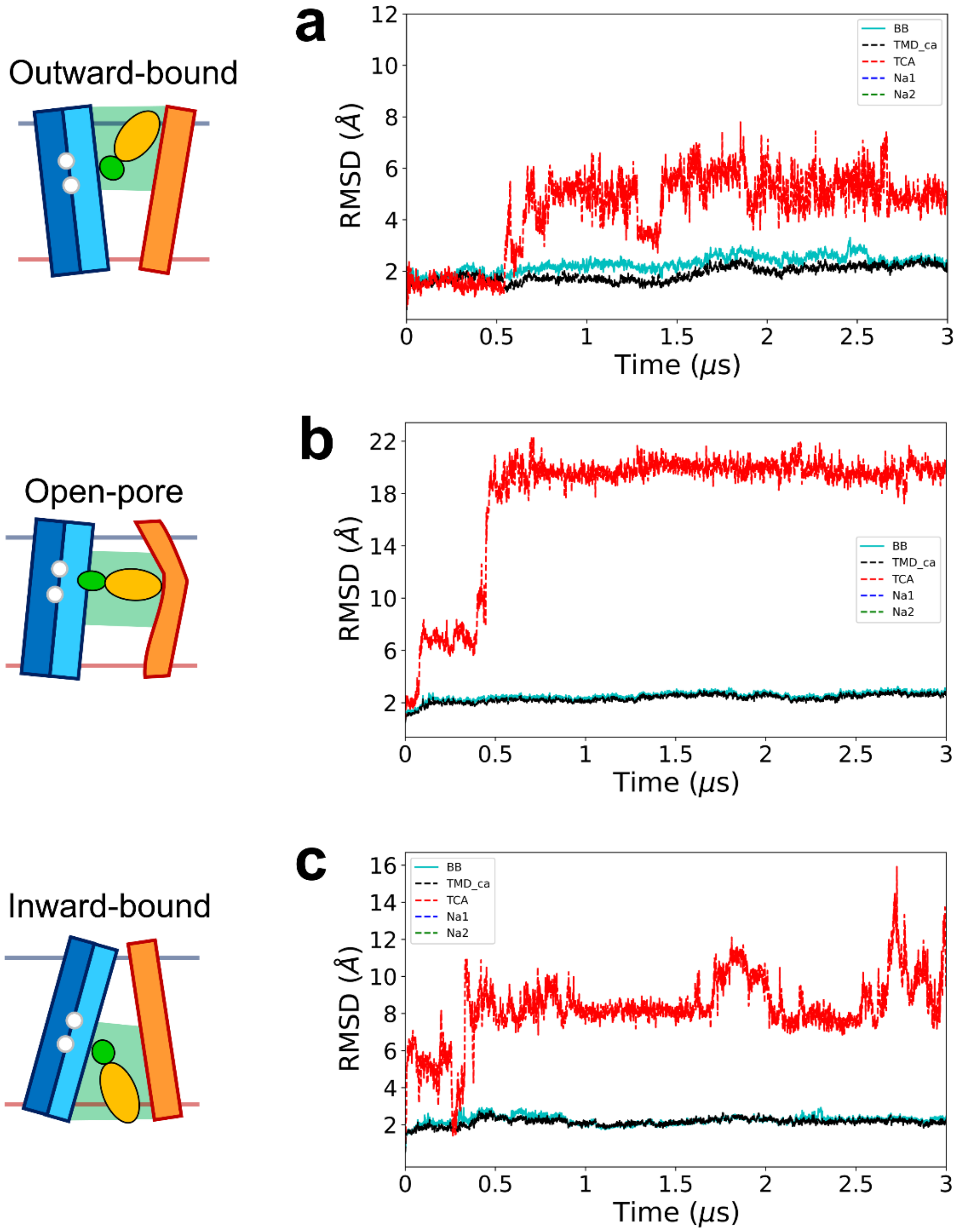

**Supplementary Fig. 12. Time evolution of RMSDs for three TCA-bound hNTCP systems without $Na^{+}$-binding in 3 μs MD simulations.** The initial conformations of TCA-bound hNTCP are represented in the left cartoon.

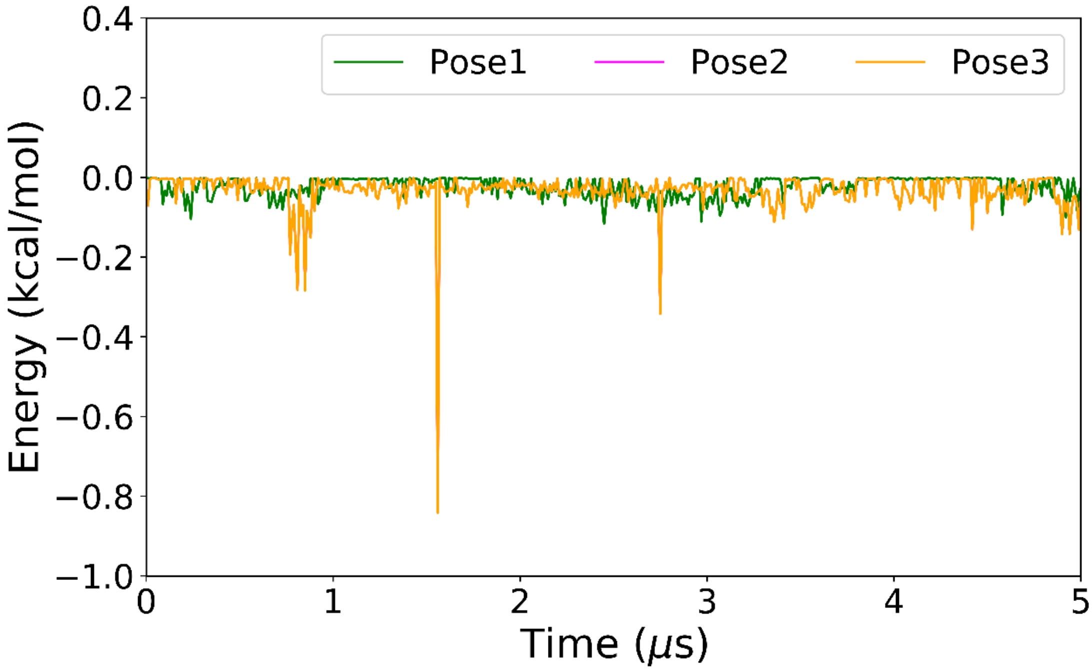

**Supplementary Fig. 13. The interaction energy between TCA and two $Na^+$ ions under three TCA-bound states in 5 μs MD simulations.**

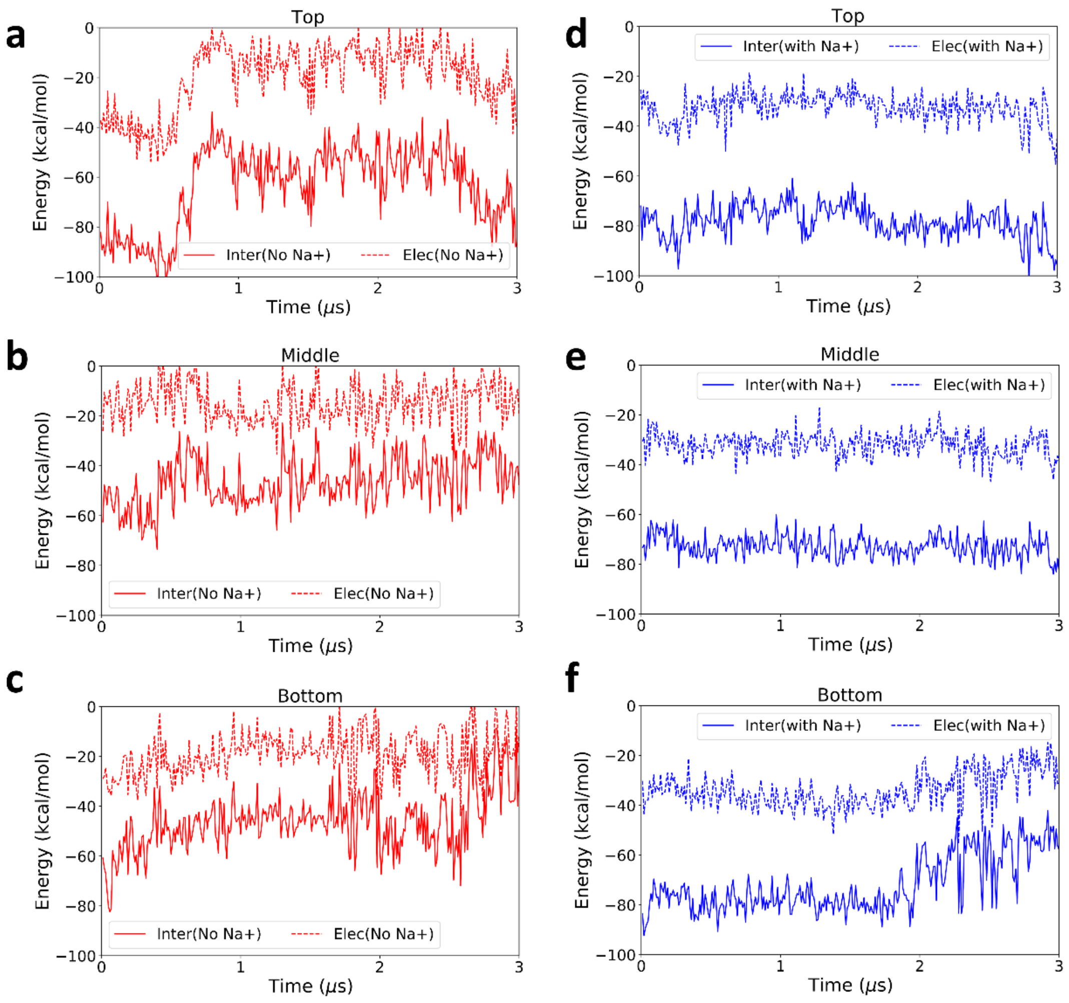

**Supplementary Fig. 14. The interaction energy between TCA and hNTCP under three TCA-bound states in 3 μs MD simulations. a-c** The three complexes without $Na^+$-binding to X-motif. **d-f** The three complexes with $Na^+$-binding to X-motif. Solid lines represent the total interaction energy between TCA and hNTCP, and dashed lines represent the contribution of electrostatic interaction between TCA and hNTCP.

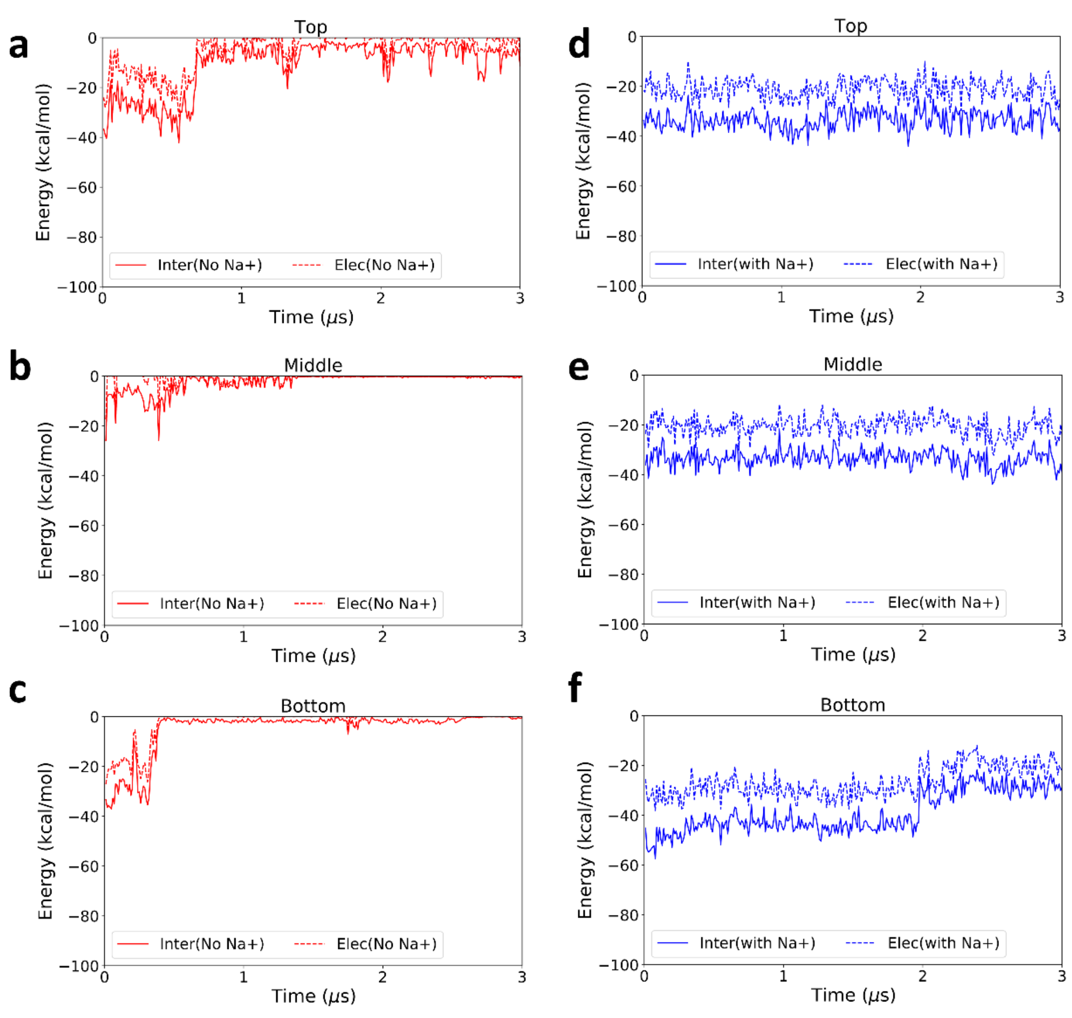

**Supplementary Fig. 15. The interaction energy between TCA and X-motif under three TCA-bound states in 3 μs MD simulations. a-c** The three complexes without $Na^+$-binding to X-motif. **d-f** The three complexes with $Na^+$-binding to X-motif. Solid lines represent the total interaction energy between TCA and X-motif, and dashed lines represent the contribution of electrostatic interaction between TCA and X-motif.

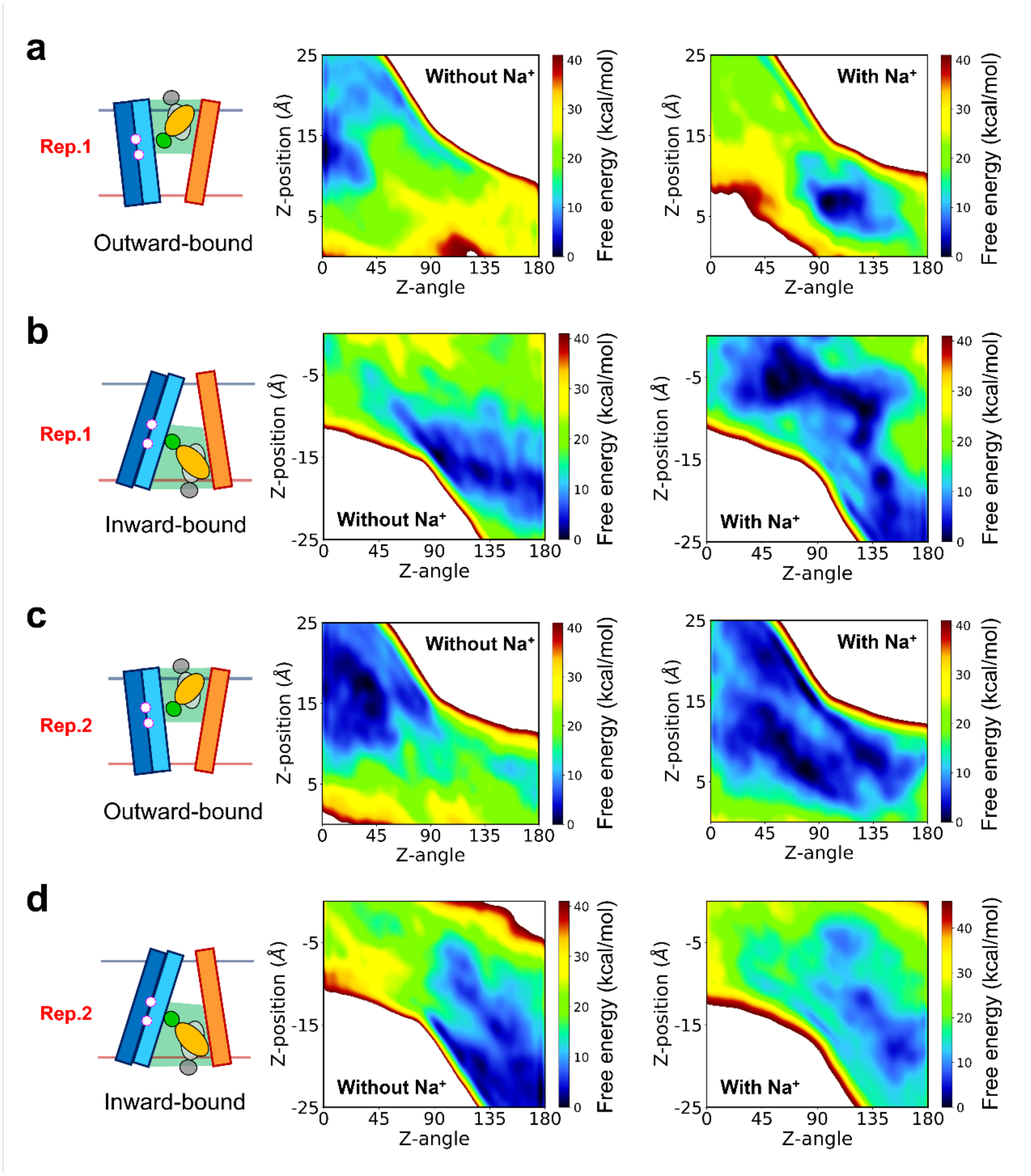

**Supplementary Fig. 16. The free energy profiles constructed from the extra two independent replicates of 2D-MetaD simulations. a-b** In the independent replicate 1, the free energy surfaces of TCA translocation in the extracellular pocket (**a**) and intracellular pocket (**b**) are reshaped by $Na^{+}$-binding states. **c-d** In the independent replicate 2, the free energy surfaces of TCA translocation in the extracellular pocket (**c**) and intracellular pocket (**d**) are reshaped by $Na^{+}$-binding states. Compared to the Rep. 0 results in Figure 6, these two additional sets of independent replicates were initialized with different equilibrated conformations and random seeds for initial velocities.

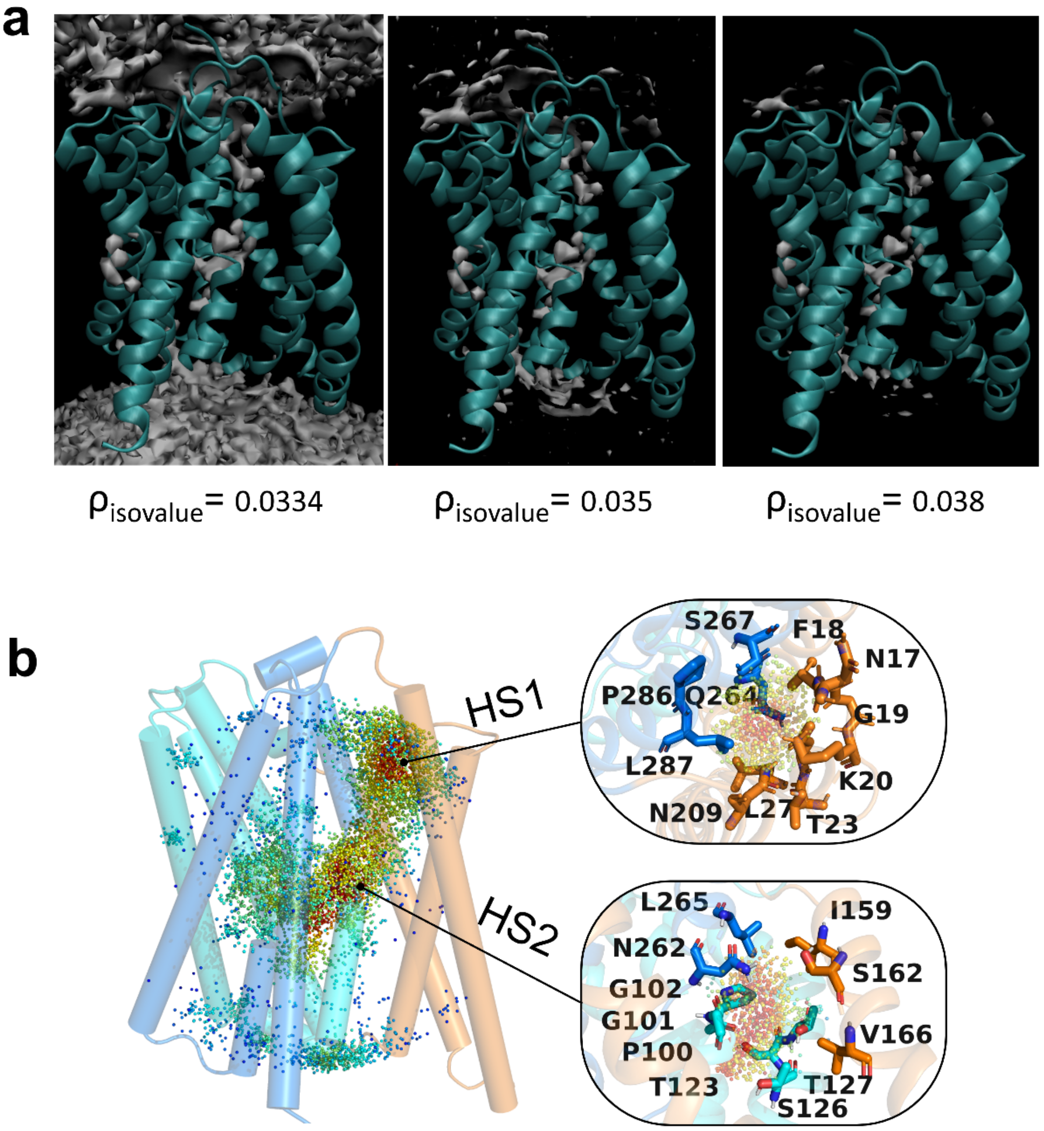

**Supplementary Fig. 17. A continuous hydration pathway that connects the X-motif and the water environment. a** Iso-surface with three iso-values of water density in 5 μs MD simulations. **b** Two hydration sites (HS1 and HS2) are identified inside hNTCP.

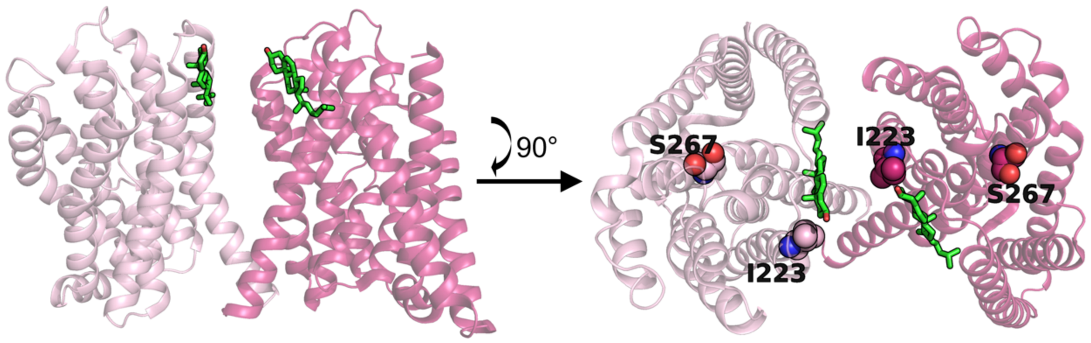

**Supplementary Fig. 18. Substrates located at the extracellular interface of the predicted dimer structure.** The monomer structure is extracted from the cryo-EM structure (PDB id: 7ZYI), which built multiple substrates in hNTCP. The dimer structure is constructed according to the work by Zeng et al, and key residues mentioned in their work to identify the dimer interface are labeled. Substrates are highlighted in green.

**Supplementary Table 1. Summary of MD simulations to understand the mechanisms of hNTCP-mediated TCA transport.**

| System ID | Protein | $Na^+$ | Substrate | Simulation | Time | Simulation purpose |
|---|---|---|---|---|---|---|
| **1** | 7pqq | 2 $Na^+$ | TCA (1) | MD | 5 µs | Dynamic investigation of TCA binding and $Na^+$ effects on TCA binding. |
| **2** | 7pqq | 2 $Na^+$ | TCA (2) | MD | 5 µs | Dynamic investigation of TCA binding and $Na^+$ effects on TCA binding. |
| **3** | 7pqg | 2 $Na^+$ | TCA (3) | MD | 5 µs | Dynamic investigation of TCA binding and $Na^+$ effects on TCA binding. |
| **4** | 7pqq (MD) | N/A | N/A | MD | 5 µs | Dynamic investigation of apo hNTCP |
| **5** | 7pqq (MD) | N/A | N/A | MD | 5 µs | Dynamic investigation of apo hNTCP |
| **6** | 7pqg (MD) | N/A | N/A | MD | 5 µs | Dynamic investigation of apo hNTCP |
| **7** | 7pqq (MD) | N/A | TCA (1) | MD | 3 µs | Dynamic investigation of TCA binding and $Na^+$ effects on TCA binding. |
| **8** | 7pqq (MD) | N/A | TCA (2) | MD | 3 µs | Dynamic investigation of TCA binding and $Na^+$ effects on TCA binding. |
| **9** | 7pqg (MD) | N/A | TCA (3) | MD | 3 µs | Dynamic investigation of TCA binding and $Na^+$ effects on TCA binding. |
| **10** | 7pqq | 2 $Na^+$ | N/A | MD | 3 µs | Whether $Na^+$-binding independent of TCA binding. |
| **11** | 7pqg | 2 $Na^+$ | N/A | MD | 3 µs | Whether $Na^+$-binding independent of TCA binding. |

| | | | | | | |
|---|---|---|---|---|---|---|
| **12** | 7pqq (MD) | 2 $Na^+$ | TCA (1) | MetaD | 100 ns | Thermodynamic investigation of $Na^+$ affects TCA translocation. |
| **13** | 7pqg (MD) | 2 $Na^+$ | TCA (3) | MetaD | 100 ns | Thermodynamic investigation of $Na^+$ affects TCA translocation. |
| **14** | 7pqq (MD) | N/A | TCA (1) | MetaD | 100 ns | Thermodynamic investigation of $Na^+$ affects TCA translocation. |
| **15** | 7pqg (MD) | N/A | TCA (3) | MetaD | 100 ns | Thermodynamic investigation of $Na^+$ affects TCA translocation. |
| **16** | 7pqq (MD) | 2 $Na^+$ | TCA (1) | MetaD | 900 ns | Thermodynamic investigation of conformational transition of hNTCP. |
| **17** | 7pqq (MD) | 2 $Na^+$ | TCA (2) | MetaD | 900 ns | Thermodynamic investigation of conformational transition of TCA-bound hNTCP. |
| **18** | 7pqq (MD) | 2 $Na^+$ | TCA (2) | MetaD | 900 ns | Thermodynamic investigation of conformational transition of TCA-bound hNTCP. |
| **19** | 7pqg (MD) | 2 $Na^+$ | TCA (3) | MetaD | 900 ns | Thermodynamic investigation of conformational transition of TCA-bound hNTCP. |

Note: TCA (1), TCA (2), and TCA(3) represent the three initial binding poses 1-3. Simulations 4-9 start from the stable conformations sampled from simulations 1-3. Simulations 12-19 start from the corresponding equilibrium conformations after ~1 μs classical MD simulation. Simulations 12-15 are performed to investigate the effect of $Na^+$-binding on substrate translocation, adding bias potential on TCA, each system has three MD replica runs. Simulations 16-19 are performed to investigate the cooperation of substrate translocation and conformational changes, adding bias potential on both TCA and hNTCP. Simulations 16-17 are set for sampling outward-facing ↔ open-pore transition, and simulations 18-19 are set for open-pore ↔ inward-facing transition.

**Supplementary Table 2. Defined CVs to characterize the conformations of TCA and hNTCP and to perform MetaD simulations.**

| CV | Description | Involved residues | Add bias |
|---|---|---|---|
| 1 | Extracellular gate size: Distance between TM1 and TM8 at the extracellular side | L25, A26, L27, S28<br><br>T268, I269, L270, N271 | No |
| 2 | Middle pore size: Distance between TM6 and TM8 at the middle position | C198, S199, V200, A201<br><br>Y289, M290, I291, F292 | No |
| 3 | Intracellular gate size: Distance between TM6 and TM9 at the intracellular side | R180, P181, Q182, Y183<br><br>W305, C306, Y307, E308 | No |
| 4 | Z-position: Position of the center of mass of heavy atoms of sulfate head in TCA | TCA | Yes |
| 5 | Z-angle: Angle of Z-direction, and vector of sterol tail to sulfate head in TCA | TCA | Yes |
| 6 | Native contact difference: Deviations in native contact refer to three TCA-bound states | Total 58 residue pairs: P22-A92, V32-G102, F33-N103, M34-L104, L35-S105, F36-N106, F37-V107, I38-F108, M39-S109, L40-L110, S41-A111, L42-M112, G43-K113, C44-G114, T45-D115, P22-V272, V23-N271, T45-R249, A207-F283, V204-P286, T203-L287, L197-Q293, I194-E296, I193-G297, G191-L299, G190-L300, K189-I301, I188-A302, V187-I303, Y186-F304, R185-W305, M184-C306, A207-V272, S206-N271, L205-L270, V204-I269, | Yes |

| | | |
|---|---|---|
| | | T203-T268, V202-S267, A201-C266, V200-L265, S199-Q264, L197-N262, I193-T258, G190-S255, K189-V254, V187-R252, Y186-R251, R185-C250, M184-R249, M192-V107, G191-F108, G190-S109, K189-L110, I188-A111, V187-M112, Y186-K113, R185-G114, M184-D115 |

**Supplementary Table 3. Parameters used in well-tempered 2D-MetaD simulations.**

| Biased CV | Width | Height (kJ/mol) | Bias-factor | Lower bound | Upper bound |
|---|---|---|---|---|---|
| Z-position | 0.1 | 5 | 50 | -25 | 25 |
| Z-angle | 4 | 5 | 50 | 0 | 180 |
| Native contact difference | 0.1 | 5 | 50 | -1 | 1 |
